# Proteome-scale tissue mapping using mass spectrometry based on label-free and multiplexed workflows

**DOI:** 10.1101/2024.03.04.583367

**Authors:** Yumi Kwon, Jongmin Woo, Fengchao Yu, Sarah M. Williams, Lye Meng Markillie, Ronald J. Moore, Ernesto S. Nakayasu, Jing Chen, Martha Campbell-Thompson, Clayton E. Mathews, Alexey I. Nesvizhskii, Wei-Jun Qian, Ying Zhu

## Abstract

Multiplexed bimolecular profiling of tissue microenvironment, or spatial omics, can provide deep insight into cellular compositions and interactions in healthy and diseased tissues. Proteome-scale tissue mapping, which aims to unbiasedly visualize all the proteins in a whole tissue section or region of interest, has attracted significant interest because it holds great potential to directly reveal diagnostic biomarkers and therapeutic targets. While many approaches are available, however, proteome mapping still exhibits significant technical challenges in both protein coverage and analytical throughput. Since many of these existing challenges are associated with mass spectrometry-based protein identification and quantification, we performed a detailed benchmarking study of three protein quantification methods for spatial proteome mapping, including label-free, TMT-MS2, and TMT-MS3. Our study indicates label-free method provided the deepest coverages of ∼3500 proteins at a spatial resolution of 50 µm and the highest quantification dynamic range, while TMT-MS2 method holds great benefit in mapping throughput at >125 pixels per day. The evaluation also indicates both label-free and TMT-MS2 provide robust protein quantifications in identifying differentially abundant proteins and spatially co-variable clusters. In the study of pancreatic islet microenvironment, we demonstrated deep proteome mapping not only enables the identification of protein markers specific to different cell types, but more importantly, it also reveals unknown or hidden protein patterns by spatial co-expression analysis.

## Introduction

Spatial omics is a rapidly evolving field that seeks to provide a deep understanding of tissue microstructures and functions by simultaneously measuring many biomolecules at high spatial resolution ^1, 2, 3, 4^. These multiplexing measurements not only reveal complex cellular and molecular compositions but also provide unique insights into cell-to-cell communications and interactions in both healthy and diseased tissues. Spatial omics has been largely fueled by various microscopic imaging and next-generation sequencing methods such as Visium spatial transcriptomics^1^, MERFISH^5^, SeqFISH^6^, Slide-Seq^7, 8^, and DBiT-seq^9^. Most of these methods were developed for the detection and quantification of transcripts with varying levels of resolution, throughput, and multiplexing capabilities. While spatial transcriptomics provide valuable insights into cellular heterogeneity, they do not directly infer the expression levels of proteins, which are the functional molecules in signaling pathways^10, 11^. Currently, the most commonly used methods for spatial proteomics are antibody-based techniques such as immunohistochemistry (IHC)^12, 13^, GeoMx ^14^, mass cytometry and MIBI-TOF, ^15, 16, 17^ and CODEX^18^. Although these immuno-affinity approaches can map protein abundances down to subcellular resolution, their multiplexity and specificity can be limited by the availability and quality of antibodies. Matrix-assisted laser desorption/ionization (MALDI) ^19^ or solvent desorption ^20^ coupled with mass spectrometry provides an alternative approach for spatial proteomics, but it is limited to detecting small molecular weight and abundant proteins due to the limitation of intact protein ionization and detection.

Mass spectrometry (MS)-based proteomics, or bottom-up proteomics has been developed for profiling proteins in various tissues, offering the deepest coverage of over 10,000 proteins per tissue sample^21^. Recent advances in mass spectrometry and sample preparation have greatly increased proteomic sensitivity, making it possible to characterize the proteome at single-cell level^22, 23, 24^. Additionally, improvements in multiplexing protein/peptide labeling and fast liquid chromatography (LC) separations further improve the throughput, allowing for large-scale proteomic analysis within reasonable instrument time^25, 26, 27^. Boosted by these advances, bottom-up proteomics has been applied to profile protein abundance on tissue sections with deep proteome coverages and high spatial resolution ^28, 29, 30, 31, 32, 33^. In most of spatial proteomics workflows, sample isolation was performed by laser microdissection (LMD or LCM). The excited tissue pixels were then digested with trypsin using a one-pot protocol for maximal recovery. For example, our group developed an LCM-nanoPOTS (nanodroplet processing in one-pot for trace samples) platform to achieve the quantitative profiling of >2000 proteins at 50-100 µm spatial resolution ^30, 31, 32, 33^. The LCM-nanoPOTS not only enables the study of cell-type or microstructure-specific proteome at single-cell level ^32, 33, 34, 35^, but also allows for unbiased spatial mapping of the whole region of interest with an imaging-like way ^30^. Deep Visual Proteomics (DVP) seamlessly integrated whole slide imaging, machine learning-based cell segmentation, laser microdissection, and TIMS-TOF mass spectrometry to perform spatial single-cell proteomics ^28, 36^. Alternatively, many non-LCM approaches such as 3D-printed micro-scaffold^29^ and Expansion Proteomics (ProteomEx) ^37^ were also demonstrated for deep spatial proteomics analysis, greatly improving the accessibility of the evolving spatial technologies.

Spatial proteome mapping or imaging of the whole tissue section or region of interest (ROI) has attracted increasing interest because of its ability to provide a comprehensive view of the tissue organization^29, 30^. In proteome mapping, the whole tissue region is pixelated by laser dissection or blade cutting, followed by sample preparation and LC-MS analysis. Typically, the protein abundances were obtained with label-free quantification. While label-free approach provides great simplicity and robustness, the throughput is inherently limited, as each isolated tissue pixel requires a single LC-MS run (e.g., 30-min)^38^. A promising way to improve the throughput is to integrate multiplexed peptide labeling with tandem mass tags (TMT), which is well demonstrated in single-cell proteomics ^22, 24^. Theoretically, using the newly released TMTpro reagent, maximal 18 tissue pixels can be multiplexed and measured within a single LC-MS run, corresponding to a throughput of >400 pixels per day if a 1-hour LC gradient is used. Despite the great potential, the use of multiplexed labeling in proteome mapping has not been evaluated. In this study, we performed a detailed benchmarking study of three protein quantification methods for spatial proteome mapping, including label-free, TMT-MS2, and TMT-MS3. To make direct comparisons, we employed serial human pancreatic sections from a healthy donor and focused on islet (endocrine) and the adjacent acinar (exocrine) regions. We compared the proteome coverages, throughput, quantification accuracy, and precision across the three methods. Our results suggest that the choice of quantification method has a great impact on the performance of proteome mapping. Each method has its own advantages and limitations, and thus project goals and experimental designs should be taken into consideration when choosing the most appropriate method.

## Results

### Study design

To benchmark the performance of label-free and multiplexed TMT labeling workflow for proteome mapping, we selected three commonly-used quantification strategies in low-input or near-single-cell samples, including label-free data-dependent acquisition with match-between-runs (LFQ-MBR, shorted as LFQ),^39^ TMT labeling with MS2 quantification (TMT-MS2),^22, 40, 41^ and TMT labeling with MS3 quantification (TMT-MS3). To minimize the biological variations from heterogeneous tissue samples, we collected pancreas tissue serial sections with a thickness of 10 µm from a healthy human donor and focused on the contrast of islet-acinar regions (**Fig. 1a)**. A spatial resolution of 50 µm with each “pixel” containing approximate 10-20 cells, was used in considering both the collection success rates and tissue heterogeneity. Pancreatic tissue consists of two major cell populations, including endocrine cells (e.g., alpha, beta, delta cells) in the islet regions and exocrine cells (e.g., acinar cells) in the acinar regions. The well-defined tissue anatomy allows us to evaluate the performance differences among these MS methods. All the sample collection and proteomic processing were performed in nanoPOTS to improve overall sensitivity and reproducibility (**Fig. 1a)**^31^.

**Fig. 1.**
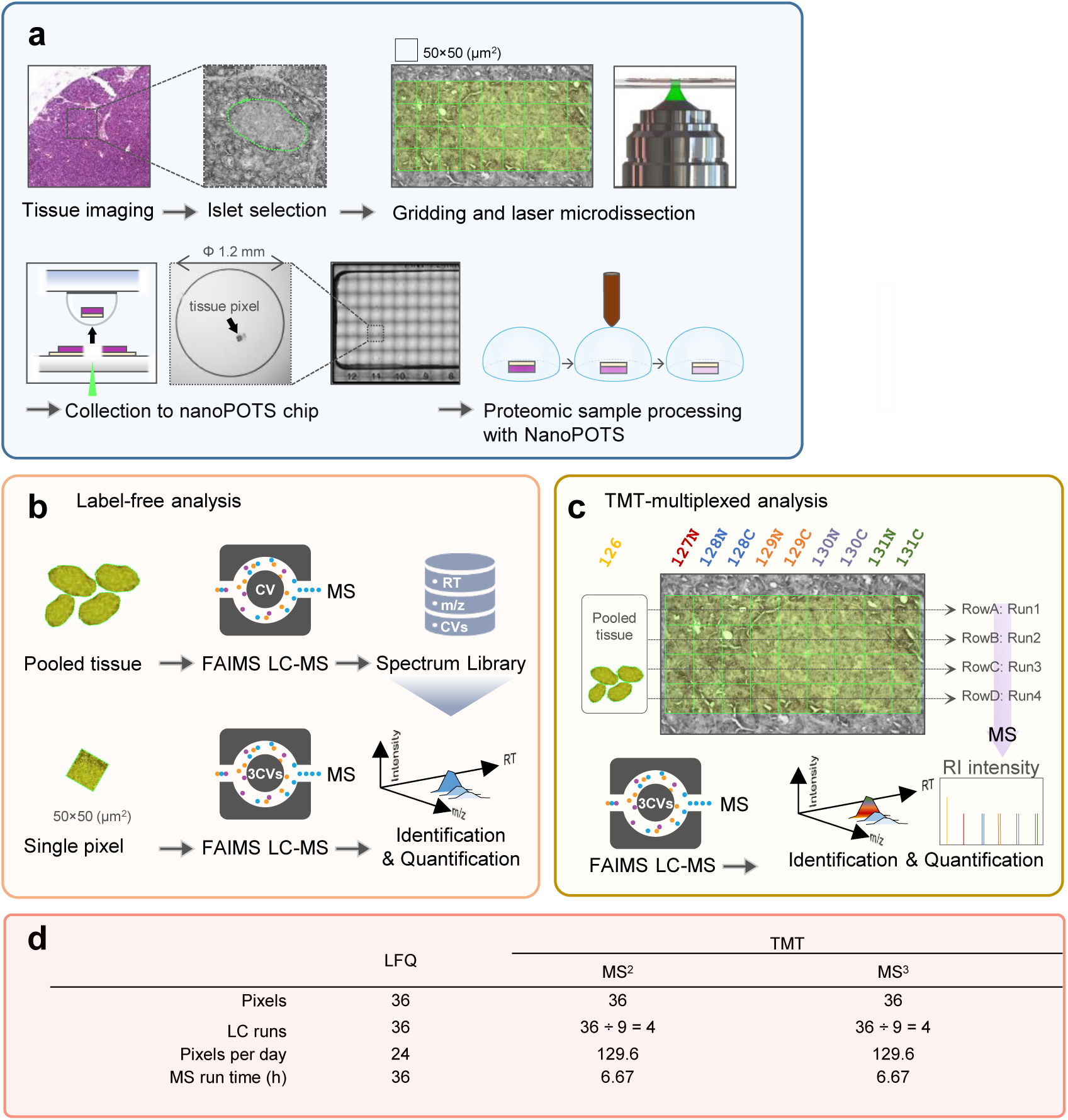
Deep proteome mapping using three different quantification approaches. (**a**). Schematic illustration of proteome mapping workflow from tissue imaging, selection of region of interest, pixelation and laser capture microdissection, sample collection to nanoPOTS chip, and proteomic sample preparation; (**b**). Workflow of TIFF-based label-free quantification (LFQ) for single pixels. High-amount pancreas tissue samples were analyzed to construct a peptide library containing LC retention time, m/z, and FAIMS CV. The peptide features in single-pixel runs were identified by matching to the peptide library based on three dimensional features; (**c**). TMT-based multiplexed analysis. TMT11 was used to label pixels in the same row together with a reference sample from a pooled tissue digest; (**d**). Tables of the total number of LC-MS runs and total run times required for 36 pixels using the three strategies.

For LFQ approach, we used a high field asymmetric waveform ion mobility spectrometry (FAIMS)-enhanced data acquisition strategy (TIFF, transferring identification based on FAIMS filtering) to increase proteome depth and reduce data missingness (**Fig. 1b**)^39^. To construct a spectral library, a bulk amount of sample from a mix of acinar and islet regions were analyzed repeatedly on LC-FAIMS-MS with three different FAIMS CVs. To match the library size used in multiplexed TMT analysis, each library sample consisted 20 ng peptides, which is ∼20 × of peptide from a single 50-µm pixel. In the previous TIFF study^39^, the samples were analyzed with a 60-min LC gradient, corresponding to a throughput of 15 samples per day. We first evaluated the impact of LC gradient times (15, 30, and 60 min) on the proteome coverage using 0.2 ng of peptides as a model sample. Reducing gradient times from 30 to 15 min significantly reduces proteome coverages by 14.8%, while no significant differences were observed when reducing gradient times from 60 min to 30 min (**Supplementary Figure 1**). Therefore, a 30 min LC gradient time was selected in this study to maximize both coverages and throughput. One concern of TIFF method is that most peptides are identified by match-between-runs (MBR), which is prone to false discovery and poor quantification. To address this issue, we employed IonQuant^42^, a tool that can control the false discovery rate (FDR) of the MBR, to analyze the data. To assess the quantification performance of TIFF method, we performed an analysis of two-proteome mixtures with known input ratios (**Supplementary Figure 2**). The TIFF method allowed the detection of ∼2348 proteins from 0.1 ng mouse sample and 630/863 proteins from 0.01/0.03 ng *S. Oneidensis* sample, by matching to a 10 ng library containing 3235 mouse proteins and 1079 *S. Oneidensis* proteins. Encouragingly, accurate and precise quantification was achieved with TIFF method, with median ratios of 0.84 and 2.55 for mouse and *S. Oneidensis* proteins, respectively. Together, these results indicated TIFF method can provide sensitive and accurate measurement of proteins for low-input samples.

For multiplexed TMT approach, an 11-plex kit was employed for isobaric labeling with a carrier or booster channel, in which 20 ng mixture peptide was labeled with TMT-126 tag and 9 pixels from one row were labeled with other tags except TMT127C (**Fig. 1c)**. Because the proteome coverage of multiplexed workflow is directly associated with the acquired and identified MS2 spectra, we employed a gradient time of 60 min to acquire sufficient MS2 spectra. The 60-min gradient time was chosen based on previous single-cell proteomics studies from various research groups^41, 43, 44, 45^, in which 60 min to 120 min gradient times were widely used. Although it is possible to obtain better proteome coverages with longer LC gradients, it should be noted that most of the additionally identified spectra were from the 20-ng carrier samples, instead of low-abundant sample channels. Thus, extended proteome coverage with long gradient times could lead to poor quantification and increased missing values. To alleviate the ratio compression problem in the isobaric labeling method, we incorporated FAIMS to pre-fractionate peptide ions into multiple populations before MS acquisition. For TMT-MS2 approach, we employed an AGC (automatic gain control) enhanced method to acquire more ions from low-input samples.^41, 46, 47^ For TMT-MS3 approach, the real-time search algorithm was enabled to improve data acquisition efficiency.

Based on these experimental design, LFQ-MBR workflow required 36 hours to complete one proteome mapping experiment containing 36 pixels and 36 LC-MS runs, while multiplexed TMT workflows only required 4 LC-MS runs and 6.67 hours. The corresponding mapping throughputs for LFQ and TMT workflows were 24 and 129.6 pixels per day, respectively (**Fig. 1d**). Thus, TMT workflows exhibited significant benefit in throughput due to sample multiplexing.

### Benchmarking the quantification strategies for spatial proteomics

To perform a pair-wise comparison between LFQ and TMT methods, we collected two pairs of islets and their surrounding regions from adjacent tissue sections (**Fig. 2a)**. Pair 1 was analyzed with LFQ-MBR and TMT-MS2 method, and pair 2 was analyzed with LFQ-MBR and TMT-MS3 method. We first assessed the proteome coverages of these methods (**Fig. 2b, Supplementary Figure 3, Supplementary Figure 4, and Supplementary Table 1-4**). For the average numbers of detected peptides and proteins across single pixels, LFQ method provided highest numbers (*n*peptide = 19348 ± 637, *n*protein = 2937 ± 71 for pair 1; and *n*peptide = 17001 ± 511, *n*protein = 2712 ± 76 for pair 2), followed by TMT-MS2 (*n*peptide = 6637 ± 342, *n*protein = 1309 ± 26) and TMT-MS3 (*n*peptide = 4715 ± 157, *n*protein = 1022 ± 18). Venn diagrams indicated 93.9% and 96.6% proteins detected by TMT methods were included in LFQ data for pair 1 and pair 2, respectively (**Fig. 2c**). When requiring valid values for all 36 pixels in each experiment, the protein numbers in LFQ method were dropped to 1935 (pair 1) and 1804 (pair 2), while the numbers in TMT methods were dropped to 977 (MS2) and 720 (MS3) (**Fig. 2d**). Together, these data demonstrate LFQ method affords deepest coverages for both detectable and quantifiable proteins.

**Fig. 2.**
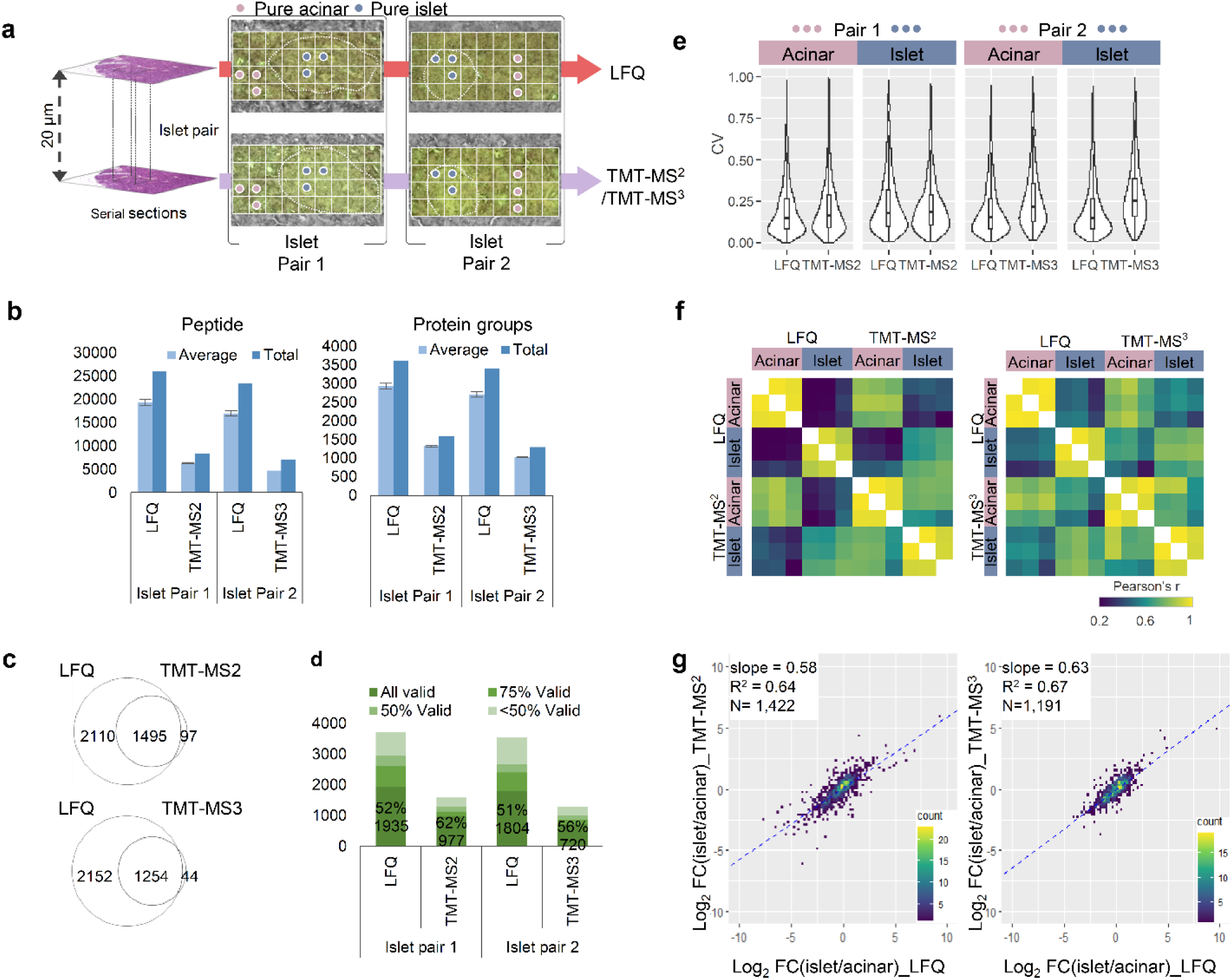
Pair-wise benchmarking of the three quantification methods. (**a**). Two pairs of pancreatic islet regions were selected from two adjacent sections. Proteome mapping of the four sections was performed with LFQ and TMT-MS2 for Pair 1, and LFQ and TMT-MS3 for Pair 2. Among a total of 36 pixels, three pixels from pure-islet (blue dots) or acinar area (pink dots) were selected for further analysis; (**b**). Average and total numbers of peptide and protein groups identified in each pixel with different quantification methods. The error bars indicate standard deviations (n = 36 for LFQ and TMT-MS2; n=35 for TMT-MS3); (**c**). Venn diagram showing the overlap between total protein identifications with different quantification methods; (**d**). Percentages and numbers of valid values across the collected pixels in the same sections at different data missing levels; (**e**). Violin plots showing the distributions of coefficient of variations (CVs) of all proteins quantified in three pure islet and acinar pixels; (**f**) Pearson correlation clustering matrix for the pure islet and acinar pixels. The color scale bar indicates the range of Pearson correlation coefficients; (**g**) Scatter plots of log2-transformed fold changes (FC) between pure islet vs acinar pixels measured in label-free and TMT-MS2 or MS3 methods. The linear regression analysis of log2FC results in regression coefficients (R^2^) and the slope value.

The use of a single database search tool (FragPipe) allows us to evaluate the identification scores and FDRs for peptides obtained from the three methods. We extracted the Probability value of each peptide, which reflects the confidence of peptide identification. As shown in Supplementary Figure 5, for the shared peptides among the three methods, we observed the highest scores in the TMT-MS2 methods, followed by the TMT-MS3 and LFQ methods. Such results are expected as both LFQ and TMT-MS3 spectra were collected from low-resolution ion trap, while TMT-MS2 spectra were from high-resolution Orbitrap. Additionally, in TMT methods, the ion population in each spectrum is summed from all TMT channels together with 20× carrier channels, which is ∼28 × higher than LFQ spectrum (Supplementary Figure 5). The ion population difference also explains the better score of TMT-MS3 method in comparison to LFQ method. For unique peptides, the Probability scores are relatively lower than shared ones, mainly due to the low abundances. Despite these differences, most of the Probability scores are > 0.99, indicating low FDRs for all these methods. We also evaluated if there are any biases in peptide identification with LFQ and TMT methods. We compared the physicochemical characteristics for peptides identified commonly and uniquely by each method, focusing on attributes such as hydrophobicity and size (**Supplementary Figure 6**). The data indicated no significant differences in molecular weight, hydrophobicity, gravy score, and peptide length across all the methods.

Next, we compared the quantitative performance of these three methods. Because the collected tissue pixels are highly heterogeneous covering both the Langerhans islet and its surrounding acinar cells, the biological variations can affect technical evaluation results. To alleviate this, we picked out pixels from “pure-acinar” and “pure-islet” regions from the two adjacent sections based on morphological annotation (**Fig. 2a**). The two distinct regions allowed us to evaluate the quantification performance such as coefficient of variation (CV) and Pearson’s correlation for LFQ and TMT methods. As shown in **Fig. 2e**, similar CV distributions with median values from 14.5% to 17.9% were observed for pair 1 experiment, indicating robust quantification is achievable with both LFQ and TMT MS2 methods. In pair 2 experiment, we observed significantly higher CVs of 25.1% for TMT MS3 data compared to 14.7% for LFQ data. We attributed the reduced measurement precision to the low sensitivity of RTS-SPS-MS3, because ion loss could occur during multiple-level ion isolation and fragmentation process.^48^ We also assessed the relationship of CVs with precursor intensities (**Supplementary Figure 7**). As expected, relatively higher CVs are observed at low precursor intensities for all the three methods. The correlation is weaker with TMT methods as reporter-ion intensities are associated with both precursor intensities and the MS/MS sampling position on the elution peak.

We performed pairwise correlation analysis across the 12 pixels in each pair using proteins with CV values higher than 15%. As expected, higher correlations were observed between the tissue pixels of the same cell types than different sample types. For LFQ method, the median correlation coefficients are from 0.91 to 0.97 for the same cell types, and from 0.24 to 0.40 for different cell types (**Fig. 2f**). For TMT methods, similar correlation coefficients from 0.95 to 0.97 are observed for same cell types. However, the correlation between different cell types (0.69 for MS2 and 0.66 for MS3) are significantly higher than that in LFQ method. Such higher correlations indicate the measured protein profiles with TMT methods exhibit more similarity between islet and acinar cells, which we attribute to ratio compression.^49^ In addition, we also performed correlation analysis between the same cell types across different quantification methods and achieved decent results with correlation coefficients from 0.68 to 0.79 (**Supplementary Figure 8**).

To evaluate the ratio compression of TMT methods, we plotted log2-transformed fold changes between islet and acinar cells using overlapped proteins in both methods. Total 1422 and 1191 proteins were included in pair 1 and pair 2 experiments (**Fig. 2g**). The slopes of linear regression between two different methods were 0.64 for pair 1 (LFQ versus TMT MS2) and 0.67 pair 2 (LFQ versus TMT MS3), respectively. Indeed, the slope values indicated that fold changes were compressed in both TMT measurements. Interestingly, although TMT-MS3 method is expected to significantly mitigate the ratio compression,^50^ our data shows its improvement is minimal in comparison to TMT-MS2 method. The diminished improvement can be largely attributed to the use of FAIMS-based ion fractionation prior to data acquisition, as well as the lower sensitivity of MS3 method.

Together, these evaluations demonstrated LFQ method is preferred for deep proteome coverage and accurate quantification, while TMT-MS2 method is a preferred method when high throughput is required.

### Unsupervised pixel clustering for characterizing tissue features

After evaluating the quantification methods from a technical perspective, we examined their performance in tissue feature characterization by comparing tissue morphological features with proteome-based data-driven classification. Unsupervised Leiden clustering identified two major clusters corresponding to acinar- and islet-dominant pixels in both LFQ and TMT methods (**Fig. 3a-b and Supplementary Figure 9**). When clustering results were annotated on LCM tissue pixels, cluster 1 corresponded to acinar areas and cluster 2 corresponded to islet areas. Uniform manifold approximation and projection (UMAP)-based dimensional reduction analysis largely recaptured the Leiden clustering results, where the two clusters were projected into two major domains. Of note, one acinar pixel was classified into islet-dominant clusters in Pair 2 experiment with TMT-MS3 quantification method. UMAP projection indicated this pixel was located at the boundary of the two main clusters (**Supplementary Figure 9**b). Overall, UMAP analysis showed that LFQ data have better classification power than TMT-based data, which we attributed to its better accuracy and relatively larger quantification dynamic range, as we observed in the above fold change analysis (**Fig. 2g**). Because LFQ data contained over twice the number of proteins compared to TMT methods, we next investigated if the improved classification power of LFQ data was simply due to its deeper proteome coverage. We generated UMAP plots using only overlapped proteins in LFQ and TMT methods (**Supplementary Figure 10**). As expected, the separation of islet and acinar samples was slightly reduced when only overlapped proteins were used for UMAP projection. Despite this, the LFQ method still exhibited better classification power than TMT.

**Fig. 3.**
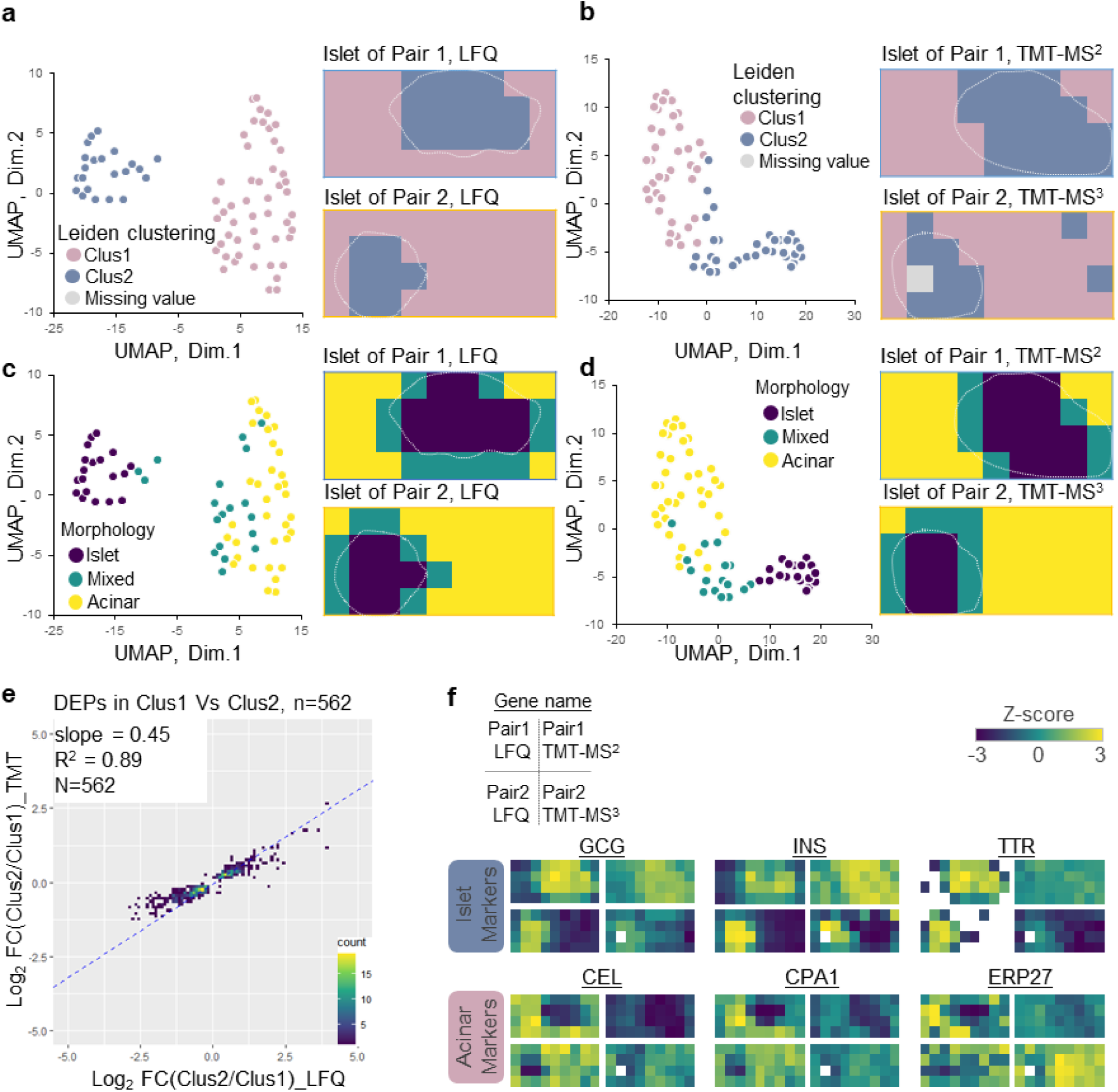
Unsupervised clustering of individual pixels. (**a-b**). UMAP pixel projection using LFQ (a) and TMT data (b) colored with Leiden clustering result (cluster 1 and cluster 2); (**c-d**) UMAP pixel projection colored with the morphological annotations (islet only, mixed, and acinar only). White dashed lines indicate the islet boundary in each section; (**e**). Scatter plots of log2-transformed fold changes for 562 differentially abundant proteins between cluster 1 vs cluster 2, which were measured in LFQ and TMT-MS2 methods. The linear regression analysis of log2FC results in regression coefficients (R^2^) and slope value; (**f**). Protein abundance maps of well-known pancreatic islet and acinar markers measured with different quantification methods.

We also evaluated the effect of missing data imputation on TMT data. Two different data filtering approaches were evaluated, including (1) >70% valid values across all pixels and (2) >70% valid values across all pixels and at least 1 valid value in each TMT batch. As shown in Supplementary Figure 11A and B, the UMAP projections were similar between non-imputed and imputed data after filtering out proteins containing <70% valid values and totally missed in at least one batch. Although the largest number of quantifiable proteins were retained when only filtering with 70% valid values, the UMAP projection appeared quite different from the other two approaches. Thus, to avoid imputation-associated artifacts, we performed imputation on proteins containing >70% valid value in all pixels and at least 1 valid value in each batch.

Based on imaging-based morphological features, we manually annotated all pixels into three cell-type groups (islet, acinar, and mix) and labeled them with different colors on UMAP plot. (**Fig. 3c-d and Supplementary Figure 9**). Pixels uniformly comprised of one type of cell were clearly separated from each other on the UMAP plot, while the pixels comprised of islet and acinar cells together were closely distributed to each other at the intermediate region. This analysis demonstrated proteome data-driven pixel classification agreed well with morphology-based annotation.

There were 562 proteins commonly identified as differentially abundant proteins (DAPs) between Cluster 1 and 2 in both methods. We calculated log2-transformed protein-abundance ratios between two clusters and examined how these ratios correlated. As we observed in the similar analysis with selected pure islet and acinar pixels (**Fig. 2g**), the slope value of 0.45 again suggested larger fold changes was obtained with LFQ method (**Fig. 3e**). Encouragingly, great linear regression coefficient of 0.89 was observed, demonstrated the two methods agreed well in quantifying the DAPs. To validate if these DAPs can explain cell populations, we used a previously reported pancreatic cell-type marker list from the human protein atlas (HPA) reference based on single-cell transcriptomics analysis ^51^. Among the 238 genes reported as acinar and islet markers in the HPA database, 62 genes were commonly identified in our proteome data set and 30 genes (11 islet markers and 19 acinar markers) showed a significant difference between Cluster 1 and 2 (**Supplementary Figure 12**a). Correlation analysis of the 30 genes showed a clear high correlation between the same cell types across all tissue pixels analyzed in this study (**Supplementary Figure 12**b).

Finally, we manually generated proteome maps using well-known islet cell markers (pro-glucagon, GCG; insulin, INS; and transthyretin, TTR) and acinar cell markers (bile salt-activated lipase, CEL; carboxypeptidase A1, CPA1; and endoplasmic reticulum resident protein 27, ERP27) (**Fig. 3f**). For both LFQ and TMT methods, distinguishable heatmaps indicating different cell-type annotations were observed for these marker proteins (**Fig. 3f**); however, the LFQ method display better contrast between the endocrine and exocrine regions based on these markers, which we ascribed to the larger quantification dynamic ranges from LFQ method.

### Detection and clustering of spatial patterns and associated proteins

One powerful capability of deep spatial omics analysis is to identify unique spatial patterns and tissue microenvironments that are invisible in morphological examination. To evaluate the performance of LFQ and TMT datasets for spatial pattern analysis, we employed the Hidden Markov Random Field model (HMRF) ^52^ algorithm embedded in the Giotto package^53^, a spatial omics tool widely employed in spatial transcriptomics. Quantitative proteome data matrixes and the corresponding spatial coordinates (x-y) of each pixel were incorporated as input of the HMRF clustering analysis (**Fig. 4a and 4b**). For both experimental pairs, the spatial clustering algorithm revealed highly correlated protein clusters (**Fig. 4c and 4d**), each of which exhibits a similar protein-abundant pattern (**Fig. 4e and 4f**).

**Fig. 4.**
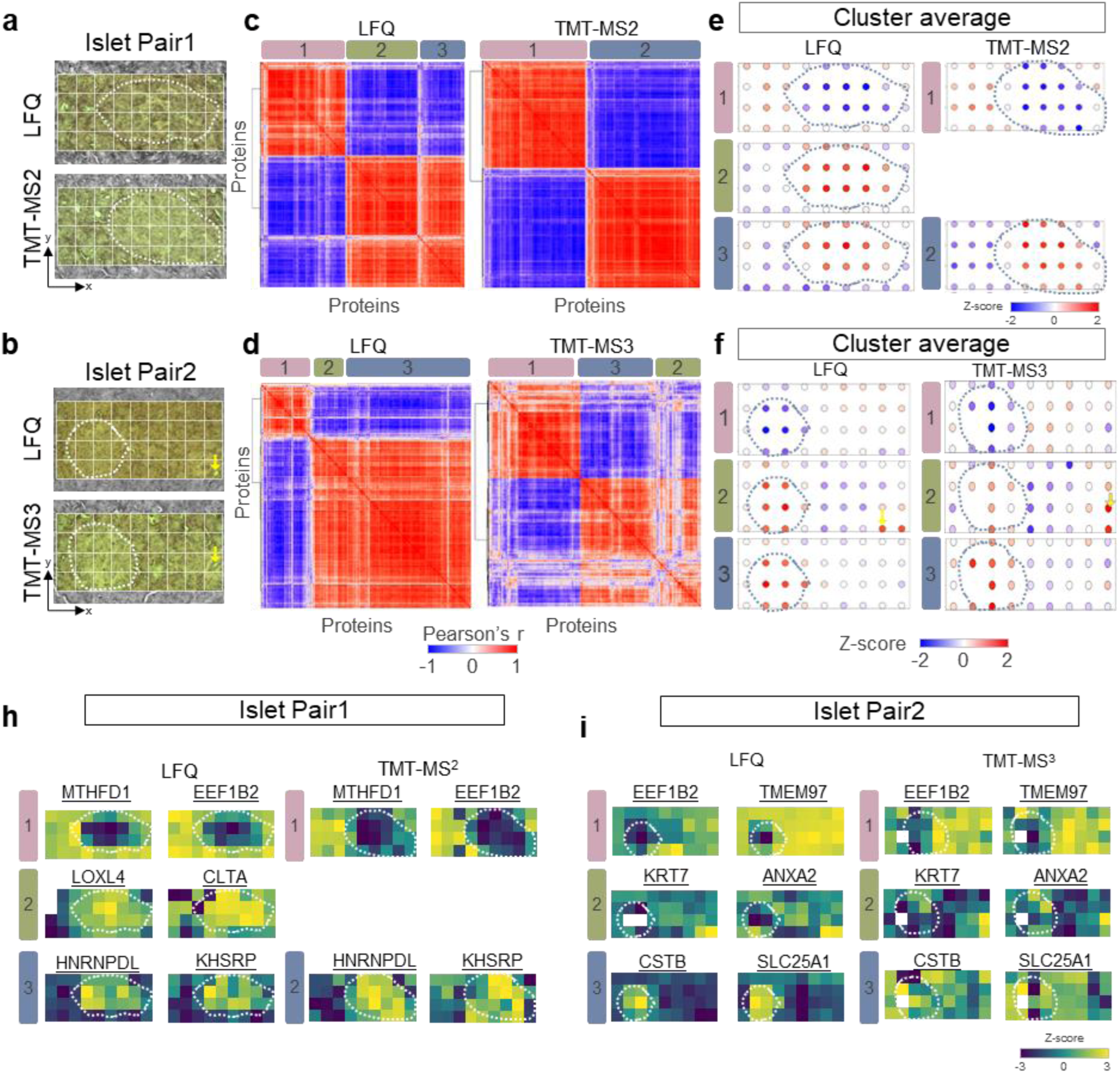
Identification of spatially co-variable proteins and clusters in pancreatic islet pairs. (**a-b**). Microscopic images of islet pairs used for each experiment. The coordinates of each pixel were determined based on the relative position from the left bottom corner. Dashed lines indicate annotated islet boundaries; (**c-d**). Correlation heatmaps showing the clustering of spatially co-variable proteins across the 36 pixels in each experiment; (**e-f**). Z-scored protein abundance maps to represent protein pattern of each cluster. (**h-i**) Protein abundance maps selected from each spatial co-variable cluster.

In Pair 1 experiment, three protein clusters emerged in LFQ workflow, while only two clusters observed in TMT-MS2 (**Fig. 4c**). Cluster 1 and 3 (and 1 and 2 in TMT) contains a list of proteins enriched in either acinar cells (acinar-high) or islet cells (islet-high), which is consistent with morphological annotation (**Fig. 4e**). For examples, MTHFD1 (Methylenetetrahydrofolate Dehydrogenase) and EEF1B2 (Eukaryotic Translation Elongation Factor 1 Beta 2) are highly enriched in acinar region, while HNRPDL (Heterogeneous nuclear ribonucleoprotein D-like) and KHSRP (KH-Type Splicing Regulatory Protein) have higher abundance in islet cells (**Fig. 4h**). Interestingly, cluster 2 in LFQ workflow revealed a list of proteins with both the islet-high pattern and relatively higher abundance in the bottom-right corner of the 4th row. However, this pattern is missed in TMT-MS2 data, which we attribute to differences in islet size between the consecutive sections. Because the islet is relatively larger in the TMT-MS2 experiment, the bottom-right corner region of the 4th row is almost occupied by the islet cells (**Fig. 4a**).

In Pair 2 experiment, both LFQ and TMT-MS3 datasets comprised three spatial patterns (**Fig. 4d**), one of which showed a unique pattern (cluster 2) other than islet-high and acinar-high patterns (**Fig. 4f**). In this cluster, proteins with the highest abundance were localized in the bottom right corner. Interestingly, these highlighted pixels could be exactly matched to a morphological feature containing a small duct with a diameter of ∼10 µm (**Fig. 4b**). These differential abundant proteins consist extracellular matrix proteins and proteins with functions of cell adhesion and structural support. Indeed, we observed highly enriched KRT7 (keratin) and ANXA2 (annexinA2) proteins in these pixels, which are well-known markers for duct cells (**Fig. 4i**). In comparison with LFQ and TMT-MS2 methods, the TMT-MS3 exhibits many less correlated proteins (**Fig. 4d**), which we attributed to the low data quality of MS3-based quantification as we observed in previous section.

Finally, we applied spatial network analysis to map another islet region (**Fig. 5**). Based on label-free quantitative mapping of 3558 proteins, we observed three distinct spatial patterns, including expected “islet high” (cluster 1), “acinar high” (cluster 3), and an unexpected “gradient expression” pattern (cluster 2) (**Fig. 5b**). Proteins in cluster 2 showed a gradient expression from top left to bottom right, regardless of cell-type difference. Cluster 1 and 3 included not only canonical marker genes shown in **Fig. 3f**, but also included additional genes that were not reported as pancreatic markers previously. Proteins in the latter included genes UCHL1 (ubiquitin carboxyl-terminal hydrolase), TIMP1 (metalloproteinase inhibitor 1), and CLU (clusterin) in cluster 1 and ZNF503 (Zinc finger protein 503), PHGDH (d-3-phophoglycerate dehydrogenase), and GSTA2 (glutathione S-transferase A2) in cluster 3 (**Fig. 5c**), indicating potential pancreatic islet or acinar markers. Total 113 proteins were included in the unexpected cluster 2, such as SRSF2 (serine/arginine-rich splicing factor 2), NCAPG (condensing complex subunit 3), and HAT1 (Histone acetyltransferase type B catalytic).

**Fig. 5.**
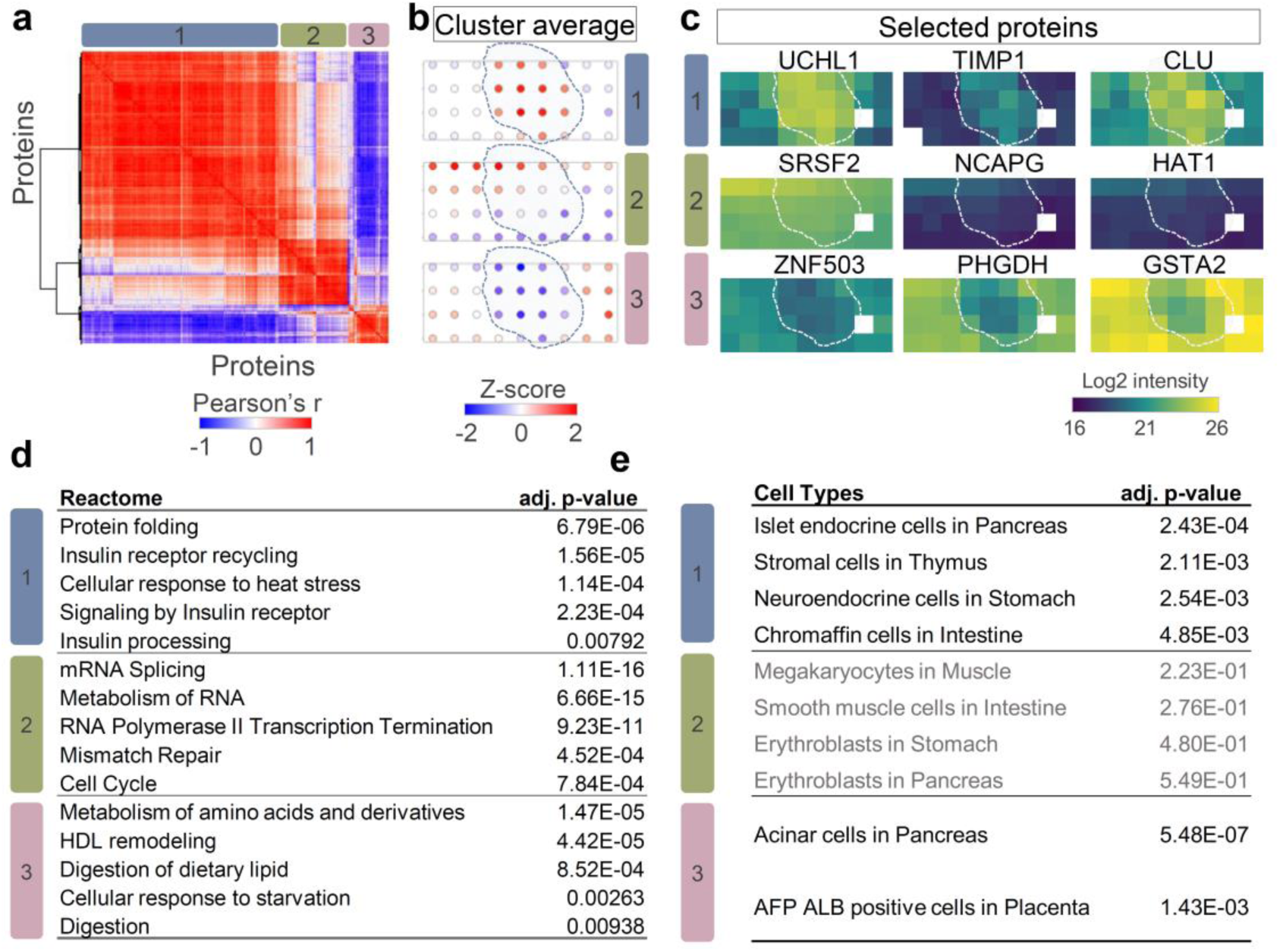
Identification of hidden tissue features and spatial patterns with spatial covariation analysis. (**a**). Correlation heatmap of three clusters containing 500 spatial co-variable proteins. Color bar indicates the range of Pearson correlation coefficients; (**b**). Z-scored protein abundance maps to represent protein pattern in each detected cluster. The dashed lines indicate islet boundaries. (**c**). Protein abundance maps selected from each spatial co-variable cluster; (**d-e**). Reactome pathway enrichment analysis (d) and cell-type enrichment analysis (**e**) using protein lists in the three clusters. Note that there is no significantly enriched cell type in cluster 2.

To understand the functional difference of the proteins with different spatial patterns, we performed Reactome pathway analysis using the three protein clusters. As shown in **Fig. 5d**, we found significant enrichment of insulin signaling pathways, such as “Insulin receptor recycling” and “Insulin processing” in cluster 1. The digestion and lipid metabolism-related pathway, such as “Metabolism of amino acids and derivatives” and “Digestion of dietary lipid” were presented in cluster 3. These function categories agree well with the major cellular compositions in islet and acinar regions. Encouragingly, we also obtained significant pathway enrichment for proteins in cluster 2. These proteins were enriched in Reactome pathways associated with cell cycling, such as “mRNA splicing”, “RNA polymerase II transcription termination”, and “cell cycle”. The pathway analysis suggested the cells in the top-left corner region were more active in protein synthesis and cell dividing, likely due to nutrition or oxygen gradient across the tissue region. In addition to pathway analysis of spatially variable proteins, we also performed cell type enrichment analysis to evaluate if the algorithm can identify the major cell types.^54^ As shown in **Fig. 5e**, the predicted cell types with the highest scores for islet-high (cluster 1) and acinar-high (cluster 3) clusters were “islet endocrine cells” and “Acinar cells”, respectively. Meanwhile, there is no significant cell type enrichment for cluster 2 proteins.

Together, we demonstrated the integration of spatial network analysis with deep proteome mapping can identify clusters of proteins with similar abundant patterns and reveal unique protein patterns that are invisible from morphological interpretation. The pathway and cell type enrichment analysis can further provide insights into the physiological functions associated with tissue microenvironment, as well as the major cellular compositions.

## Discussion

Although proteomic quantification methods have been extensively evaluated for global proteomics, phosphoproteomics, or bulk-scale tissue proteomics ^55, 56, 57^, there is a lack of study focusing on evaluating their performance on high-resolution spatial proteomics, where the samples are both low-input and highly heterogeneous in nature. For example, the total protein amount in each tissue pixel of 50 µm square is only ∼2 ng, which is >100 × lower in bulk-scale benchmarking studies. Thus, highly sensitive MS parameters, including FAIMS pre-filtering and high injection time should be used to accommodate the detection of low ion populations. Also, in spatial proteomics, each sample/pixel is a single biological replicate and is analyzed by a single LC-MS analysis without technical replicates. Because of highly heterogeneous nature, a large number of spatial samples are usually required to detect rare events as well as to generate sufficient statistical power. High throughput analysis with short LC gradient and non-fractionated multiplexed TMT sample preparation is critical. Due to these significant differences, previous bulk-scale benchmarking studies are not suitable to guide spatial proteomics experiments.

In this study, we systematically evaluated high sensitivity and high throughput proteomics methods for proteome-scale spatial mapping of tissue sections, including the commonly-used label-free and isobaric labeling methods. Our study indicated label-free method provided the deepest coverages of ∼3500 proteins at a spatial resolution of 50 µm, corresponding to ∼10-20 cells. Quantitative comparison also revealed label-free method exhibited high dynamic ranges with large fold changes between tissue regions containing two different cell populations. Despite these merits, label-free method has no multiplexing capabilities, limiting their application in large number of pixels when instrument time is limited. Such limitation could be partially addressed by shortening gradient time^58, 59^ or employing a dual-trap or dual-column system to improve the overall throughput.^27, 60^ One important consideration when analyzing a large number of spatial samples is injection-order randomization, where pixels from different spatial locations should be injected randomly to minimize the LC-MS-associated batch effect.

On the other hand, TMT-based isobaric labeling methods offered the highest throughput of >125 pixels per day by multiplexed data acquisition of over 9 pixels, making it particularly useful for large-scale experiments. Although TMT method suffered from moderate proteome coverages and compromised protein quantification accuracy in comparison with label-free method, our analysis demonstrated it provided great agreement in term of detecting differentially abundant proteins and clustering pixels, as well as identifying spatially variable proteins. Our analysis also indicates that TMT MS3 method with RTS^61, 62^ didn’t significantly improve the quantification accuracy, largely due to reduced measurement sensitivity. Thus, TMT labeling with FAIMS pre-fractionation and MS2-based quantification is recommended. We envision the throughput and coverage can be further improved by employing high-multiplicity TMT tags (e.g., 18plex or higher) and advanced mass spectrometer with faster scanning speed (e.g., Astral Orbitrap). However, due to inherent limitations of TMT-based isobaric labeling approach such as precursor co-isolation and batch effect (Supplementary Figure 13), the approach may not be suitable for mapping subtle proteome alternations. Additional validation experiments should be performed to obtain solid biological conclusions derived from the spatial proteomics results. We employed Combat algorithm to correct the LC-MS-related batch effect. It should be noted that Combat assumes the overall sample compositions are the same across batches. In this study, we labeled each “row” with a TMT plex and incorporated both islet and acinar cells in each plex to ensure similar sample compositions. Such a design could mask the systematic difference in the “column” direction, as indicated in Figure 5. Therefore, both cell type and spatial randomization should be incorporated when applying the TMT method to map large-sized tissue samples. Alternatively, recent studies demonstrated non-isobaric labeling coupled with data-independent acquisition (DIA) offered a promising solution to boost both throughput and quantification accuracy, albeit the practical multiplicity is only 3 at present ^63, 64^.

Together, we summarized the technical characteristics of the LFQ and TMT methods in **Supplementary Figure 14**. We also provided our recommendation of the method of choice based on the project goals including coverages and pixel numbers. We suggest choosing LFQ method for low throughput applications with <200 pixels, as well as applications requiring deep proteome coverages. We also note LFQ method offers the simplest sample preparation procedures as no chemical labeling step is included. TMT-MS2-based multiplexed method naturally fits in applications requiring mapping large tissue areas or multiple serial tissue sections containing a large number of pixels.

Deep proteome mapping of human tissue holds significant potential in understanding tissue functions in health and disease states by combining traditional histological imaging with unbiased proteomic profiling. We demonstrated the LFQ approach allowed to create proteome maps of healthy human islet regions covering > 3000 proteins. We also demonstrated deep and spatially-resolved proteome mapping not only enabled to identify protein markers specific to different cell types, more importantly, it also revealed unknown or hidden protein patterns by spatial co-expression pattern analysis. As an example, we demonstrated vasculature-associated pixels and nutrition/oxygen-induced proteome gradient patterns could be discovered by a spatial clustering algorithm (e.g., HMRF). These capabilities allow us to study heterogeneity and unknown molecular patterns of human tissues, especially in diseased tissues such as type 1 diabetes and cancers. The direct access to the proteome of diseased tissues could also provide potential biomarkers or drug targets for precise diagnosis and therapeutic intervention.

## Methods

### Preparation of human pancreas tissue

Human pancreas tissue was obtained from a 17-year-old male donor. The donor was selected based on the eligibility criteria established by the HuBMAP consortium, under IRB201600029 (https://www.protocols.io/view/donor-eligibility-criteria-and-pancreas-recovery-f-b7nfrmbn), and following the ethical standards of the Declaration of Helsinki. Organ recovery and tissue processing were performed at the University of Florida following the standard protocol (https://www.protocols.io/view/human-pancreas-processing-b7gxrjxn). Tissue block embedded in CMC media was frozen with dry ice/isopentane bath and stored in -80 °C freezer.

### NanoPOTS chip fabrication

Nanowell chips were fabricated on glass slides as described previously^32^. Briefly, an array of 4 × 11 nanowells with a diameter 1.2 mm and a center-to-center spacing of 4.5 mm were fabricated on glass slides with pre-coated chromium and photoresist (25 mm × 75 mm, Telic company, Valencia, USA) based on standard photolithography and wet etching. After wet etching, the remaining glass surfaces were treated with 2% (v/v) heptadecafluoro-1,1,2,2-tetrahydrodecyldimethylchlorosilane in 2,2,4-trimethylpentane. After removing the remaining chromium layer, an array of spots with hydrophilic surface served as nanowells for tissue collection and proteomic sample processing.

### Laser capture microdissection and sample collection into nanoPOTS

First, 10-µm thickness tissue sections collected on PEN membrane slides (Carl Zeiss) were stained using Hematoxylin and Eosin staining kit (H&E staining kit, Abcam) following the manufacturer’s instruction. Before sample collection, 200-nL DMSO was pre-loaded onto each nanowell as sample capture liquid.^31^ Laser capture microdissection (LCM) was performed on a PALM MicroBeam LCM system (Carl Zeiss MicroImaging, Munich, Germany). Pixelation of tissue section was performed by drawing a gridline around the region of interest on the tissue, followed by tissue cutting and catapulting. Pancreas tissue was cut at an energy level of 40 using the “CenterRoboLPC” function of PalmRobo software. For sample collection, an energy level of delta 15 and a focus level of delta 5 were used to catapult tissue pixels into nanowells. The collected tissue sample in this study is a three-dimensional-form voxel, however, LCM is a technology for isolating two-dimensional tissue area from a thin layer, rather than a cube. Thus, we called each collected sample as a “pixel”. After collection, the nanowell chip was heated to 70 °C for 20 min to evaporate the DMSO droplet under hanging drop mode^33^. The dried chip was imaged with a microscope to confirm successful tissue collection and stored at -20 °C until use.

### Experimental Design and Statistical Rationale

Our overall experimental design is depicted in Figure 1 and Figure 2a. We generated a total of four spatial proteome maps, each consisting of a pancreatic islet and its surrounding acinar cells. These maps include 36 pixels created by dissecting a 4 × 9 grid from the tissue and collecting the samples directly into corresponding wells in a nanoPOTS chip. These spatial maps were derived from two pairs of islets, with each pair being selected from consecutive tissue sections. For each pair of islets, one islet was analyzed using LFQ, while the other was analyzed using TMT (either TMT-MS2 or TMS-MS3). This resulted in a comprehensive analysis of four islets in total (2 islets per section × 2 sections = 4 islets), allowing us to examine and compare the spatial proteomic profiles in a consistent manner between consecutive tissue sections within the same biological context. To compare the proteome difference between islet and acinar regions, 9-23 islet pixels and 13-27 acinar pixels in each section were analyzed to generate sufficient statistical power.

### NanoPOTS-based proteomic sample processing

A home-built nanoliter robotic liquid-handling platform was employed to dispense reagents into nanowells.^32, 65^ First, 200 nL of cell lysis buffer containing 0.1% (v/v) n-dodecyl-ß D-maltoside (DDM), 1 mM tris(2-carboxyethyl)phosphine (TCEP) and 0.1 M 4-(2-hydroxyethyl)-1-piperazineethanesulfonic acid (HEPES, pH 8.0) was applied into each nanowell and incubated at 70 °C for 1 h. Next, 50 nL of 10 mM of

2-chloroacetamide (CAA) in 0.1 M HEPES (pH 8.0) was added to each nanowell and incubated for 30 min at room temperature. After alkylation, protein digestion was performed overnight by addition of 50 nL of 0.01 ng nL^-1^ Lys-C (MS grade, Promega, Madison, USA) and 50 nL of 0.04 ng nL^-1^ trypsin (Promega) in 0.1 M HEPES buffer and incubated at 37 °C for 10 h. During all incubation step, the chip was placed in an upside-down direction to increase the protein extraction and digestion efficiency^33^.

For LFQ samples, the digestion was quenched by dispensing 50 nL of 5% formic acid (FA) into each well and incubated for 15 min. Samples in nanowell chip were dried out in a vacuum desiccator and stored in -20 °C until analysis.

For multiplexed labeling samples, TMT reactions were processed without any desalting steps. TMT11 plex reagents were resuspended in DMSO at a concentration of 10 ng/nL and 100 nL of each TMT tag (1 µg) were applied into each nanowell following our experimental design and incubated at RT for 1h. Next, the labeling reaction was quenched by adding 50 nL of 2.5% (w/v) hydroxylamine and incubated at RT for 15 min. All samples were then pooled together to microPOTS wells, acidified with 5% FA, dried out in a vacuum desiccator, and stored in -20 °C until analysis.

### LC-MS/MS analysis

A homebuilt nanoPOTS autosampler was employed to automatically perform sample collection, cleanup, and LC separation^23^. The LC separation system consisted of an in-house packed solid-phase extraction (SPE) column (100 µm i.d., 4 cm, packed with 5 µm, C18 packing material (300 Å pore size; Phenomenex, Torrance, CA, USA) and an in-house packed LC column (50 μm i.d., 25 cm-long, packed with 1.7 μm, C18 packing material (BEH 130 Å C18 material, Waters) heated at 50 °C with a column heater (Analytical Sales and Services Inc., Flanders, NJ, USA).

Dried peptide samples on chips were dissolved with Buffer A (0.1 % FA in water), then injected to the SPE column for 5 min with a loading buffer containing 2% acetonitrile. After washing, samples were eluted and separated at 100 nL/min using gradient of Buffer B (0.1% FA in acetonitrile). All the samples were analyzed by an LC-MS system, which consisted of an UltiMate 3000 RSLCnano System (Thermo Fisher Scientific) and an Orbitrap Eclipse Tribrid MS (Thermo Fisher Scientific) with a FAIMSpro interface.

For the LFQ analysis, a 30-minute and 100 nL/min linear gradient from 8% to 22% Buffer B (0.1% formic acid in acetronitrile) followed 9-minutes linear gradient from 22% to 35% mobile phase B was used. Peptides were ionized by applying a voltage 2400 V at the electrospray source.

For construction of spectral library, the ionized peptides were fractionated by the FAIMSpro interface with each LC-MS run utilizing a discrete FAIMS compensation voltage (CV, -45, -60, or -75 V). Fractionated ions were collected into an ion transfer tube heated at 250 °C and ions with mass range 350-1600 m/z were scanned at 120,000 resolution with an ion injection time (IT) of 118 ms and an automatic gain control (AGC) target of 1E6. The selected precursor ions with +2 to +6 charges and >1E4 intensities were isolated with a window of 1.4 m/z and fragmented by a 30% level of high energy dissociation (HCD). Fragmented peptide ions were scanned in ion trap with maximum injection time of 86 ms and AGC target of 2E4. The cycle time was set to 2 sec.

For LFQ single pixel analysis, the ionized peptides were fractionated by the FAIMSpro interface using three FAIMS CVs (-45, -60 and -75V) for each LC-MS analysis. Fractionated ions with mass range 350-1600 m/z were scanned at 120,000 resolution with an IT of 246 ms and an AGC target of 1E6. The selected precursor ions with +2 to +6 charges and >1E4 intensities were isolated with a window of 1.4 m/z and fragmented by a 30% level of HCD. Fragmented peptide ions were scanned in ion trap with maximum injection time of 86 ms and AGC target of 2E4. The cycle time in each CV experiment was set to 0.8 sec.

For multiplexed labeling analysis, a 60-minutes linear gradient from 8% to 28% Buffer B, followed by 15- min gradient to 45% at a flow rate of 100 nL/min were used. Peptides were ionized by applying voltage 2400 V at the electrospray source and ionized peptides were fractionated by the FAIMSpro interface using cycling two FAIMS CVs (-45 and -65 V).

For TMT-MS2 analysis, fractionated ions with mass range 450-1800 m/z were scanned at 120,000 resolution with an IT of 118 ms and an automatic gain control (AGC) target of 1E6. The selected precursor ions with +2 to +6 charges and >1E4 intensities were isolated with a window of 0.7 m/z and fragmented by a 35% level of HCD. Fragmented peptide ions were scanned in an Orbitrap with a maximum injection time of 150 ms and AGC target of 5E5. The cycle time in each CV experiment was set to 1.5 sec.

For TMT-MS3 analysis, fractionated ions with mass range 450-1800 m/z were scanned at 120,000 resolution with an IT of 118 ms and an automatic gain control (AGC) target of 1E6. Selected precursor ions with +2 to +6 charges and >1E4 intensities were isolated with a window of 0.7 m/z and fragmented by a 35% level of collision induced dissociation (CID). Fragmented peptide ions were scanned in ion trap with maximum injection time of 80 ms and AGC target of 1E4. For MS3 analyses, synchronized precursor selection (SPS) coupled with real-time search (RTS) strategy was applied. Real-time search was configured with a full enzyme specificity, a max variable modification of 2, a max missed cleavage of 1, and a max search time of 100 ms. Fixed modifications included TMT6plex on n-terminal and lysine residue (229.1629 Da) and carbamidomethylation on cysteine residue (57.0215 Da). Oxidation on methionine (15.9949 Da) was set as dynamic modification. Scoring threshold was set to 1.4 Xcorr, 0.1 dCn, and 10 ppm precursor tolerance. After selection of 10 notches, fragmented ions were fragmented by a 45% level of HCD. The generated ions were scanned in Orbitrap with the range of 100-150 m/z at 60,000 resolution. The MS3 maximum IT was 250 ms and AGC target was 1E5. The cycle time in each CV experiment was set to 1.5 sec.

### MS data analysis

All spectrum raw files were processed using FragPipe (v 17.1) powered by MSFragger (v. 3.5) ^66, 67^ search engine, Philosopher (ver. 4.2.1)^68^ for FDR filtering and reporting, and IonQuant^42^ for label-free MBR and quantification, and TMT-Integrator for TMT-based quantification. All mass spectrum was searched against a UniProt human (Homo sapiens) data base (Apr2, 2022 released) containing 20,318 protein sequences, 116 common contaminant sequences, and decoy protein sequences.

For LFQ-MBR analysis, following search parameters were used for MS/MS spectrum search: full tryptic specificity up to two missed cleavage sites, carbamidomethylation (57.0214 Da) on cysteine as a fixed modification, and methionine oxidation (+15.9949) and protein N-terminal acetylation (42.0105 Da) as variable modifications. The initial precursor and fragment mass tolerances were set to 20 ppm. After mass calibration, MSFragger adjusted the tolerances automatically. The identification results were filtered with 1% PSM- and protein-level FDR. For quantification, match between runs algorithm in IonQuant was activated with a matching RT tolerance of 0.4 min, m/z tolerance of 10 ppm, and an ion-level FDR threshold of 5%. The normalization was enabled, and the MaxLFQ algorithm was used to generate the protein-level intensities from the ion-level intensities.

For multiplexed labeling data, following search parameters were used for MS/MS spectrum search: full tryptic specificity up to two missed cleavage sites, TMT6plex (229.1629 Da) on lysine residue and carbamidomethylation (57.0214 Da) on cysteine as fixed modifications. Methionine oxidation (15.9949 Da), protein N-terminal acetylation (42.0105 Da), and TMT6plex (229.1629 Da) on peptide N-terminal as variable modifications. Other settings are the same as the above LFQ-MBR analysis. For quantification, TMT-Integrator was used to normalize and generate the results. The mass tolerance for TMT reporter ion was set to 20 ppm. The abundances were grouped at protein level and reference samples in TMT-126 channel were used for normalization between different batches.

We followed general approaches in single-cell or low-input proteomics studies and kept proteins identified from a single unique peptides for downstream analysis^41, 43, 44, 45^. We provided a table (Supplementary Table 5-7) containing these single-peptide protein lists for each experiment and associated them with FragPipe output table available through MassIVE repository (MSV000091531). FragPipe PDV could be used to view the annotated spectra from single-peptide proteins.

### Spatial proteomics data analysis

Statistical analyses were performed using R (v 4.2.1) and Perseus (v 1.6.15.0)^69^. For the label-free outputs, the columns of protein intensities were extracted from the report files “combined_protein.tsv” and log2 transformed. For the TMT outputs, protein intensities were extracted from the report files “abundance_protein_GN.tsv”, which contain log2 transformed and globally-normalized protein abundance obtained from sample/reference ratios and reference channel intensities. All the results shown in Figure 2 utilized log2 transformed values from the output table without any missing value imputation or batch effect correction. For proteome mapping and visualization in Figure 3-5, proteins containing >70% valid values in each tissue section (36 pixels) were considered quantifiable and kept for downstream analysis. For TMT data, proteins missing in at least one TMT batch were also filtered out. Missing values imputation based on the standard distribution of the valid values (width:0.3 and downshift: 1.8) was performed in Perseus. For TMT datasets where each map contains four TMT plexes, we performed additional batch correction based on the SVA Combat algorithm^70^..

All spatial omics analysis using Giotto package with default embedded parameters ^53^. The raw protein expression matrix from Fragpipe together with the corresponding spatial location were used to create a Giotto object. After normalization, we performed Leiden clustering and overlaid on UMAP plot. Spatially variable genes were detected using the “detectSpatialCorgenes” function and clustered using “clusterSpatialCorgenes” function in Giotto package.

The Reactome pathway analysis and cell-type enrichment analysis were conducted on the Enrichr tool (https://maayanlab.cloud/Enrichr/)^71, 72^. Briefly, the proteins included in each cluster were submitted to the Enrichr and the results of Reactome pathways and cell types were exported.

## Data availability

The mass spectrometry raw data can be accessed on the ProteomXchange Consortium via the MassIVE partner repository with the data set identifier MSV000091531 and are available at ftp://MSV000091531@massive.ucsd.edu. (User name: MSV000091531, password: Nano4441)

## Acknowledgments

We thank Dr. Matthew Monroe for his assistance in depositing the raw proteomic data into MassIVE. This work was supported by a National Institutes of Health (NIH) Common Fund, Human Bimolecular Atlas Program (HuBMAP) grant U54DK127823 (W.J.Q.). A portion of this research was also performed on a project award (60201) from the Environmental Molecular Sciences Laboratory, a DOE Office of Science User Facility sponsored by the Biological and Environmental Research program under Contract No. DE-AC05-76RL01830.

