## Supplementary information for "Proteome-scale tissue mapping using mass spectrometry based on label-free and multiplexed workflows"

**Table of Contents**

Supplementary Figure 1. Optimization of gradient time for TIFF method..	3

Supplementary Figure 2. Evaluation of quantification performance of TIFF method using two-species proteome.	4

Supplementary Figure 3. Number of peptides and protein groups in single pixels in label free approach.	5

Supplementary Figure 4. Number of peptide and protein groups in single pixels using TMT.	6

Supplementary Figure 5. Boxplots showing the distribution of peptide identification probabilities and protein intensities.	7

Supplementary Figure 6. Boxplots showing the distribution of peptide physicochemical properties	8

Supplementary Figure 7. Density plots depict the correlation between coefficients of variation (CVs) and protein intensities.	9

Supplementary Figure 8. Pearson correlation matrix for the pure islet and acinar pixels.	10

Supplementary Figure 9. UMAP plot for each islet pair and quantification methods	11

Supplementary Figure 10. Impact of proteome coverages on classification power.	12

Supplementary Figure 11. The effect of missing value imputation in the TMT datasets.	13

Supplementary Figure 12. Comparison of transcriptomic markers identified in human pancreatic tissue and the proteome markers	14

Supplementary Figure 13. Batch effect in TMT dataset and the effectiveness of batch effect correction..	15

Supplementary Figure 14. Summary of quantification methods for high-resolution spatial proteomics study.	16

**Supplementary Figure 1. Optimization of gradient time for TIFF method.** (**a**) Number of identified peptides and (**b**) proteins at three LC gradient times (15, 30 and 60 min). A model sample of 0.2 ng S. Oneidensis peptide was used.

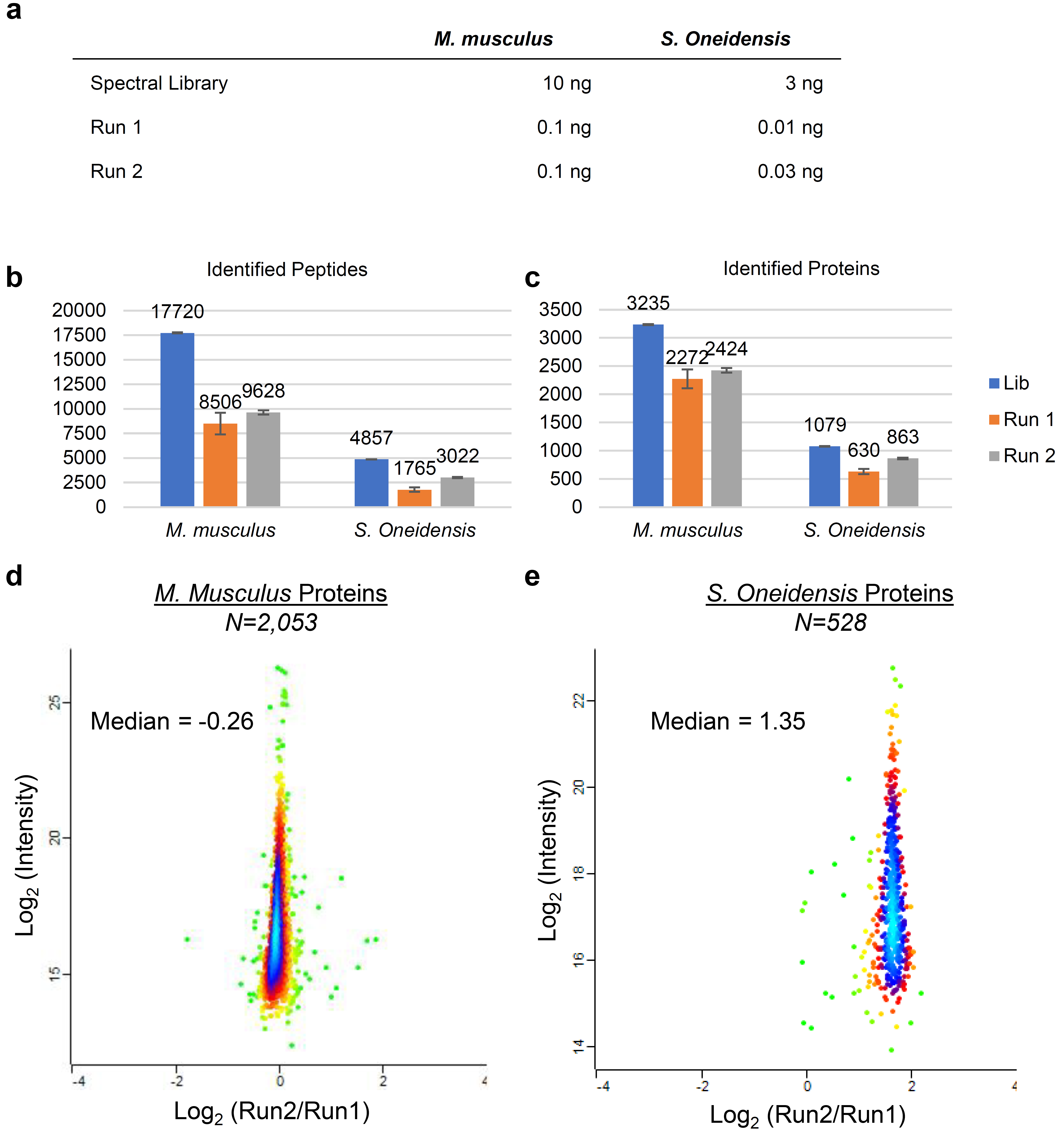

**Supplementary Figure 2. Evaluation of quantification performance of TIFF method using two-species proteome.** (**a**) Experimental design of the two-proteome study including the spectral library, Run 1, and Run 2. (**b**) Peptides and (**c**) proteins identified from two proteome mixture in library, run 1 and run 2, respectively. (**d**) Scatter plots showing protein fold changes between Run2 and Run1 for mouse proteome and (**e**) Shewanella proteome.

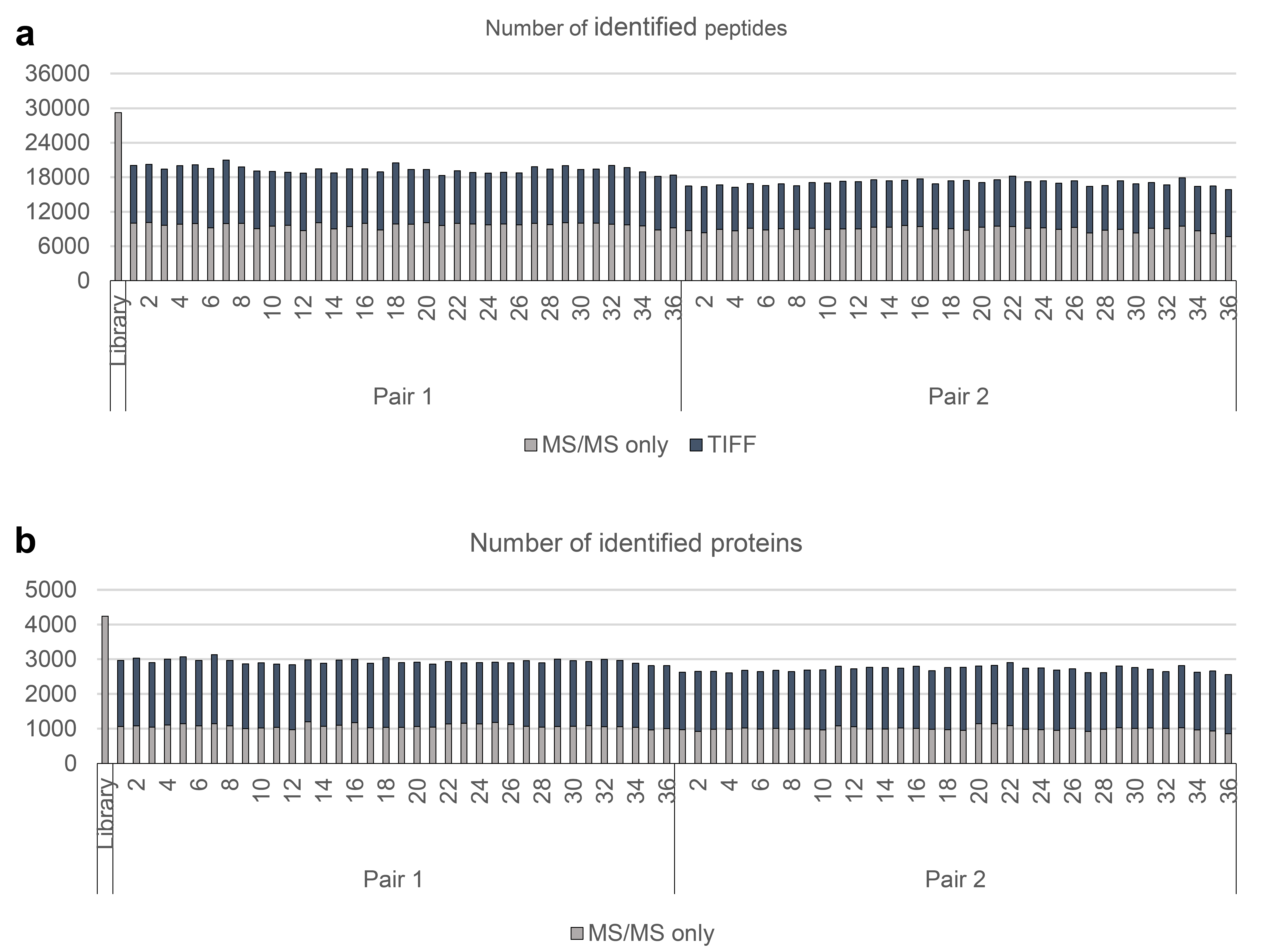

**Supplementary Figure 3.** Number of peptides (**a**) and protein groups (**b**) in single pixels identified by MS/MS only (gray) and by the TIFF method (blue) in label free approach.

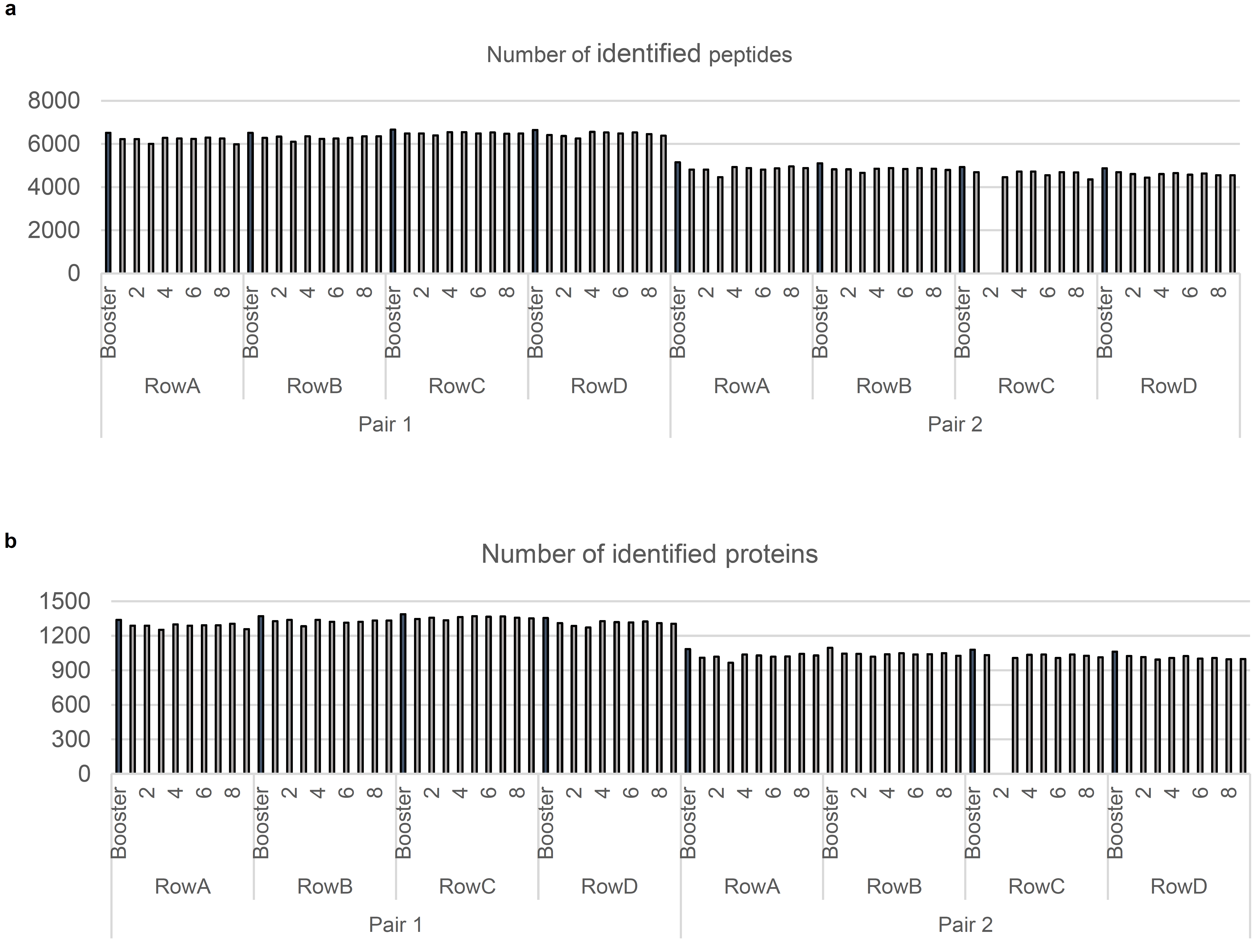

**Supplementary Figure 4.** Number of peptide (**a**) and protein groups (**b**) in single pixels using multiplexed TMT approaches**.**

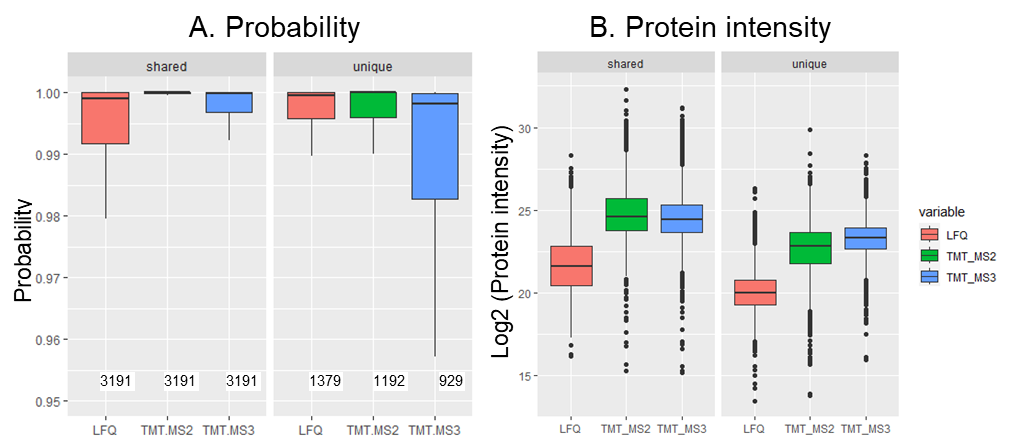
**Supplementary Figure 5.** Boxplots showing the distribution of peptide identification probabilities (A) and protein intensities (B) among different quantification methods (e.g., LFQ, TMT-MS2, TMT-MS3). Peptides shared across all methods and those uniquely identified by each method are shown.

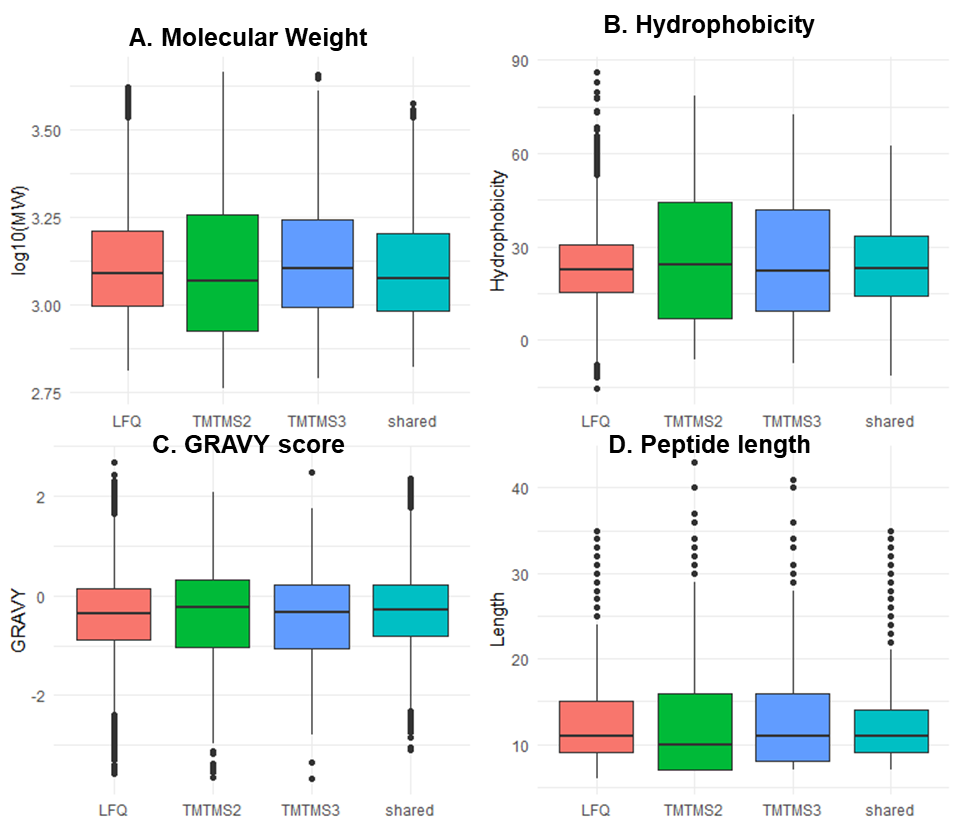

**Supplementary Figure 6.** Boxplots showing the distribution of peptide physicochemical properties in different quantification methods including (A) molecular weight, (B) hydrophobicity, (C) GRAVY score, and (D) peptide length.

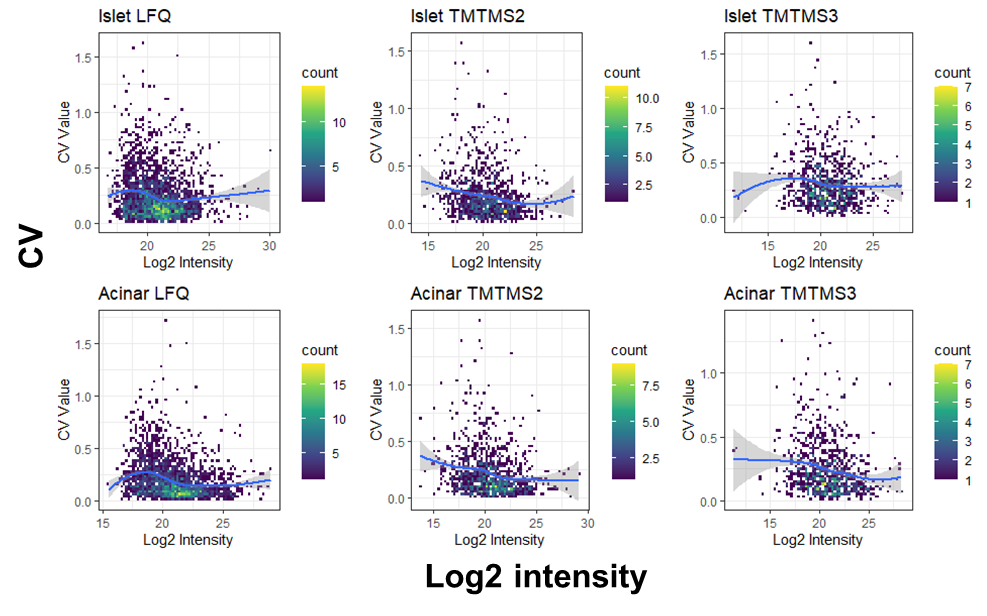

**Supplementary Figure 7.** Density plots depict the correlation between coefficients of variation (CVs) and protein intensities in different quantification methods and tissue types (islet or acinar) using the data in the main Figure 2E.

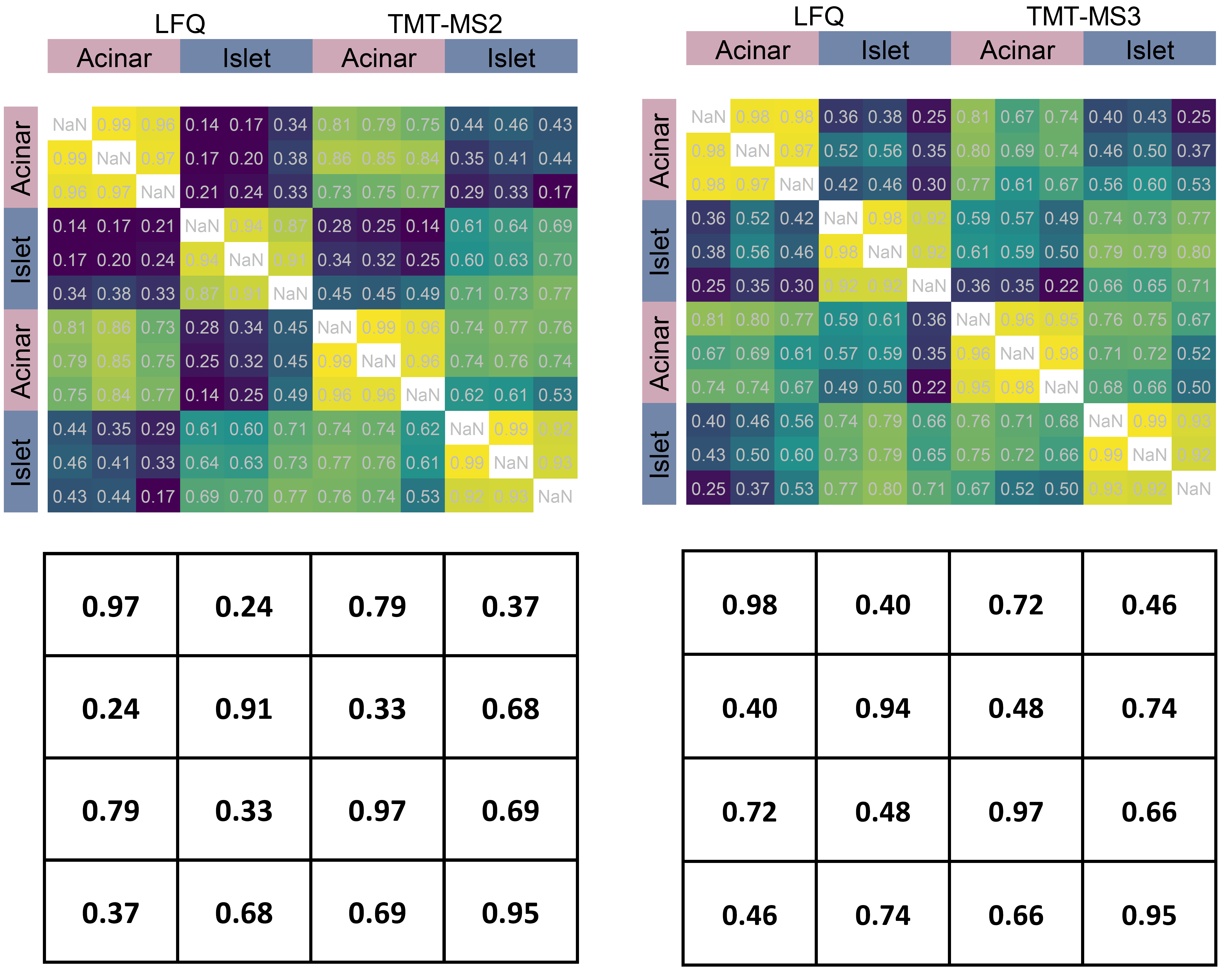

**Supplementary Figure 8. Pearson correlation matrix for the pure islet and acinar pixels.** Correlation of protein abundances between cell-types and quantification methods are shown in the correlation matrix. Median values of correlation coefficients for each cell type and quantification method were shown in the bottom panel.

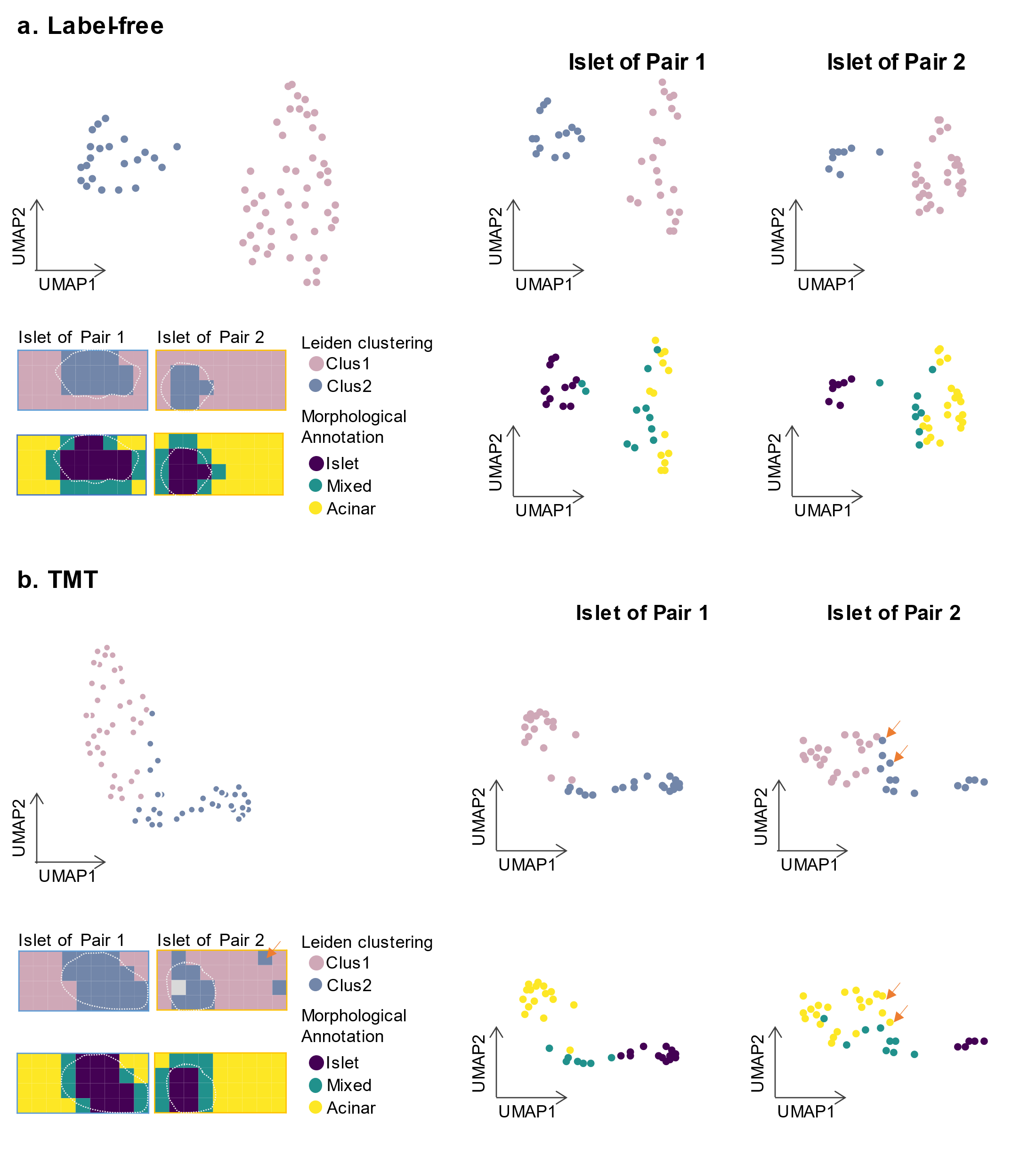

**Supplementary Figure 9.** UMAP plot for each islet pair and quantification methods for (a) label free and (b) TMT methods.

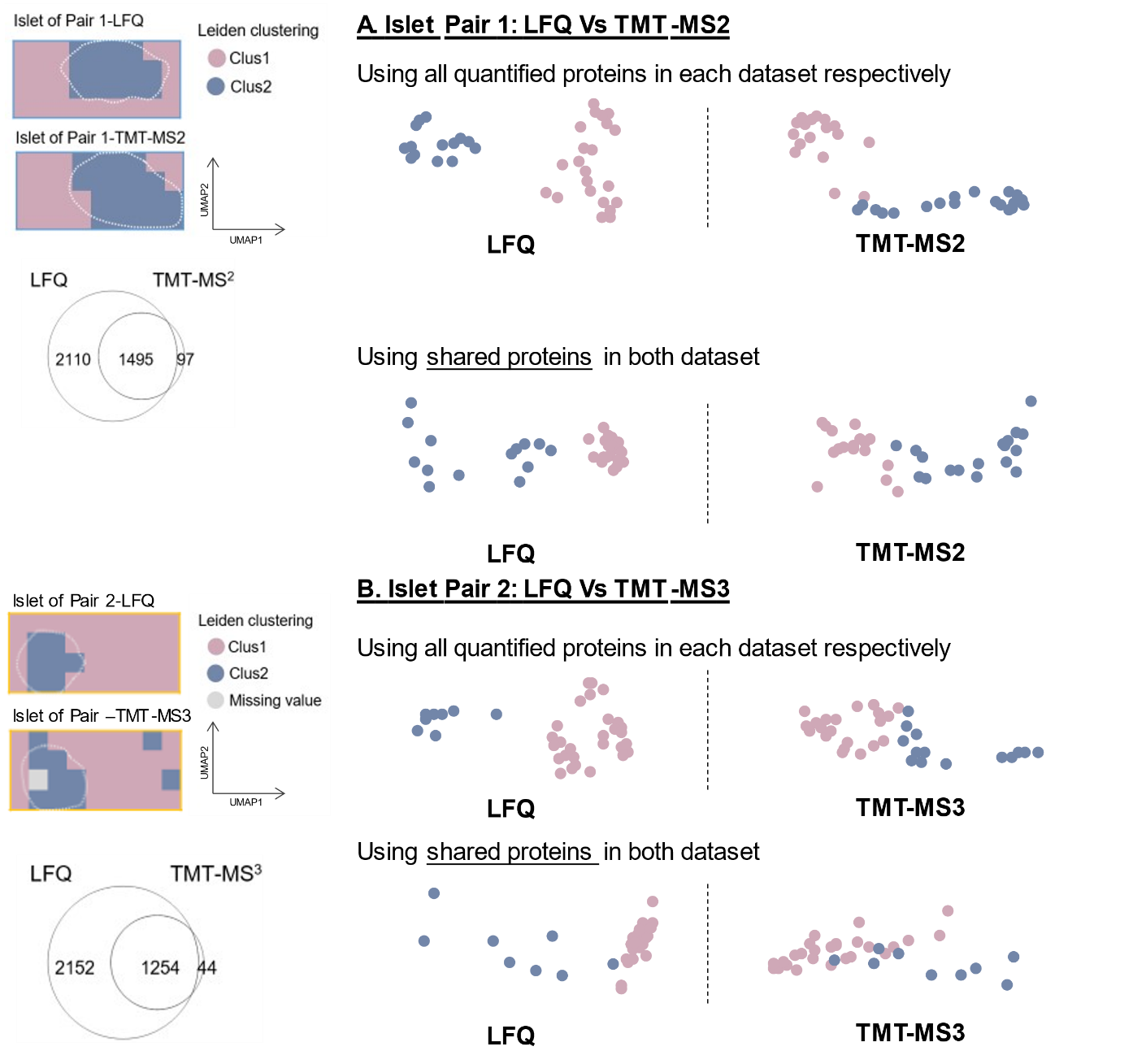

**Supplementary Figure 10. Impact of proteome coverages on classification power**. A. UMAP projection for LFQ and TMT-MS2 datasets using all quantified proteins in each dataset versus using only shared proteins. B. UMAP projection for LFQ and TMT-MS3 datasets using all quantified proteins in each dataset versus using only shared proteins. Protein numbers were indicated in Venn diagrams.

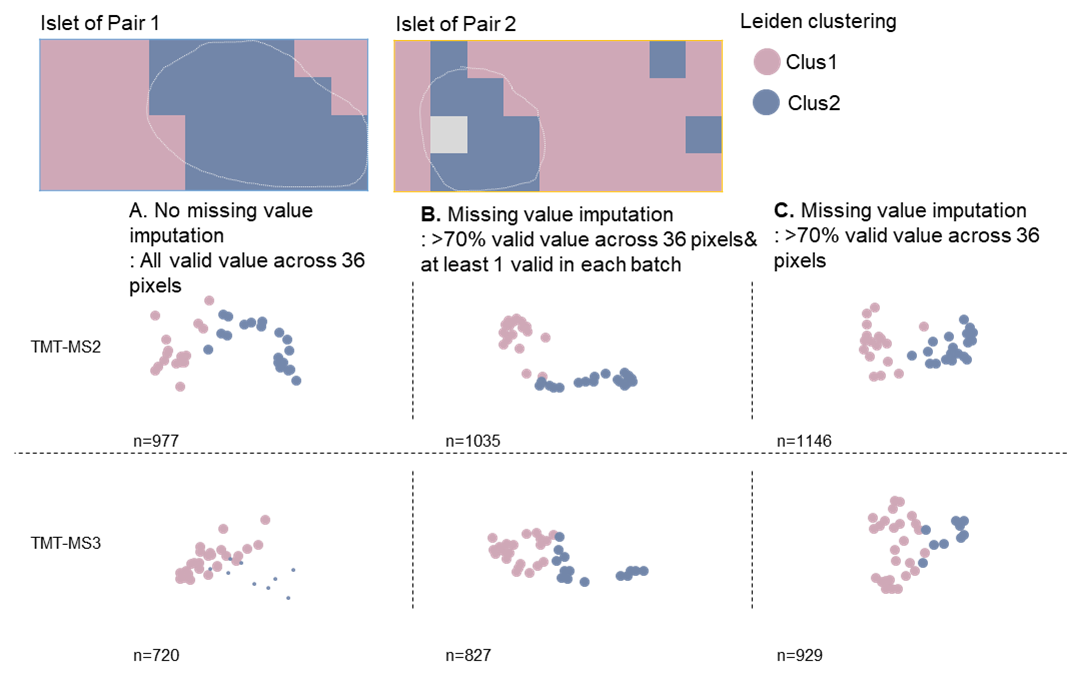

**Supplementary Figure 11. The effect of missing value imputation in the TMT datasets.** UMAP projections were performed (A) using valid values across 36 pixels without any imputation; (B) imputating the data after excluding proteins with less than 70% valid values across 36 pixels and less than 1 valid value in each TMT batch; and (C) imputating the data after excluding proteins with less than 70% valid values.

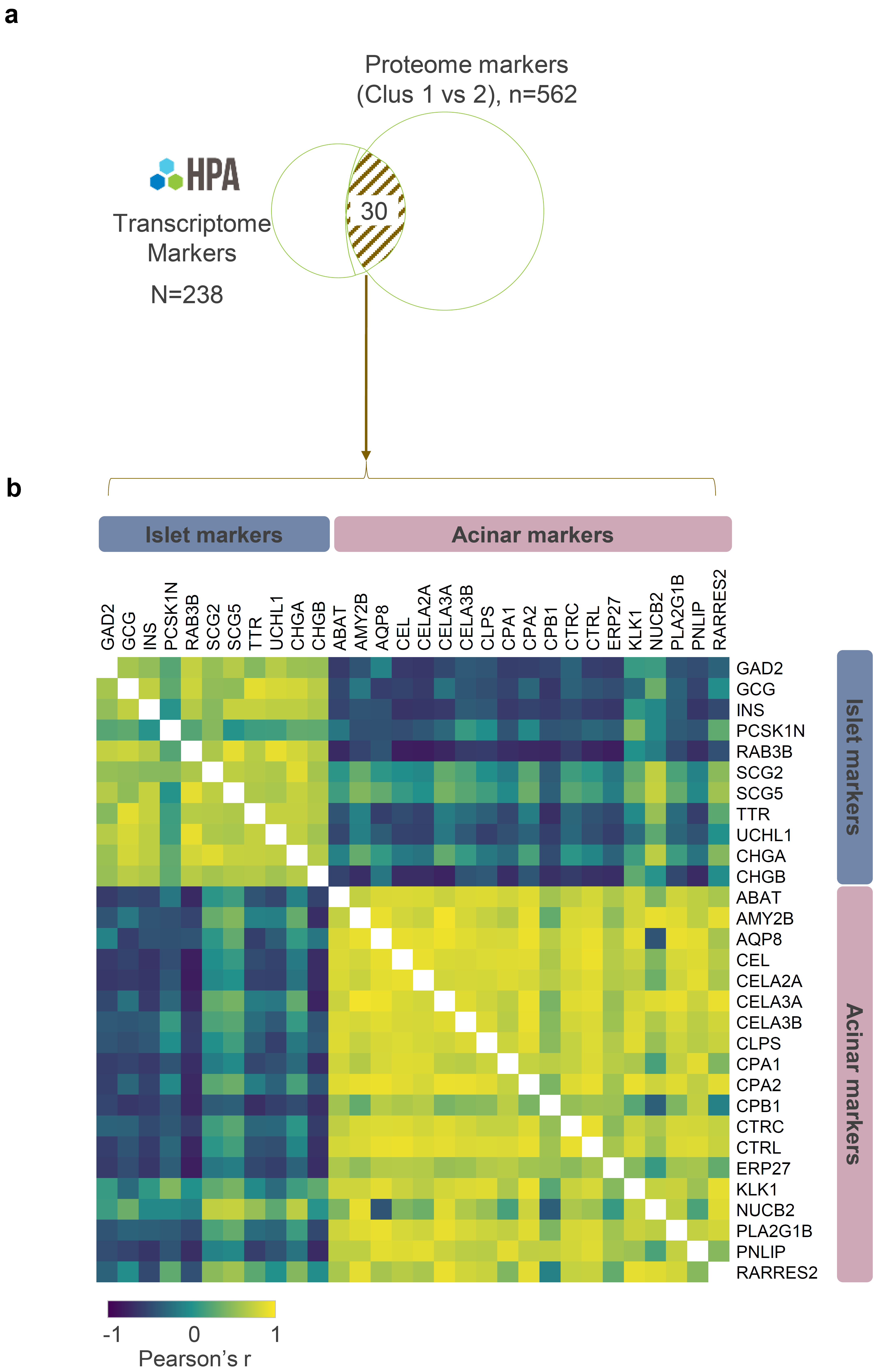

**Supplementary Figure 12.** (**a**) Venn diagram indicating the overlap of transcript markers and protein markers identified from Human Protein Atlas (HPA) database and this study, respectively. (**b**) Pearson correlation of commonly identified pancreatic markers.

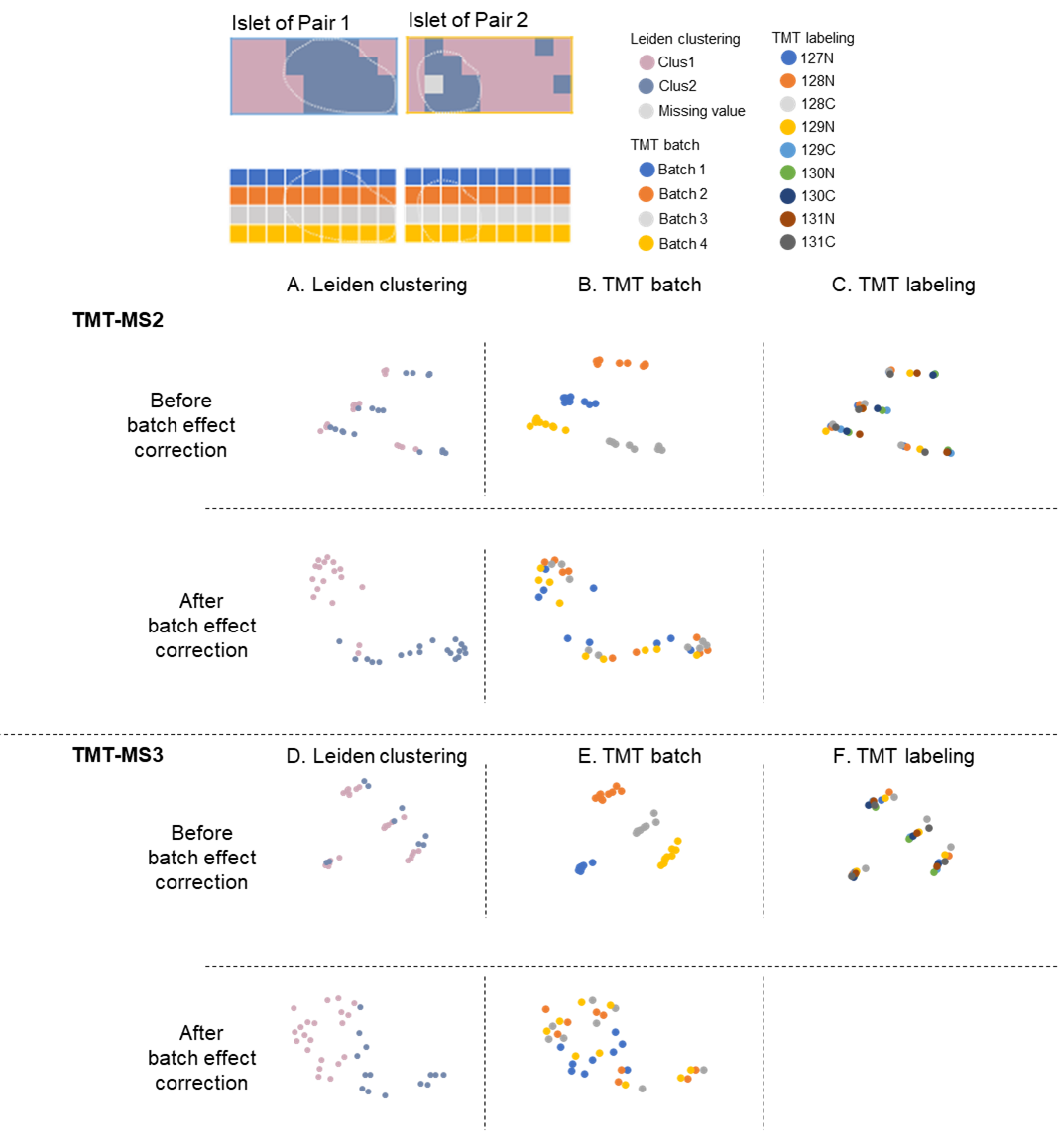

**Supplementary Figure 13. Batch effect in TMT dataset and the effectiveness of batch effect correction.** Colored UMAP plots indicate TMT batch and channel effect in TMT-MS2 (A-C) and TMT-MS3 (D-F) datasets, as well as the effectiveness of batch effect corrections.

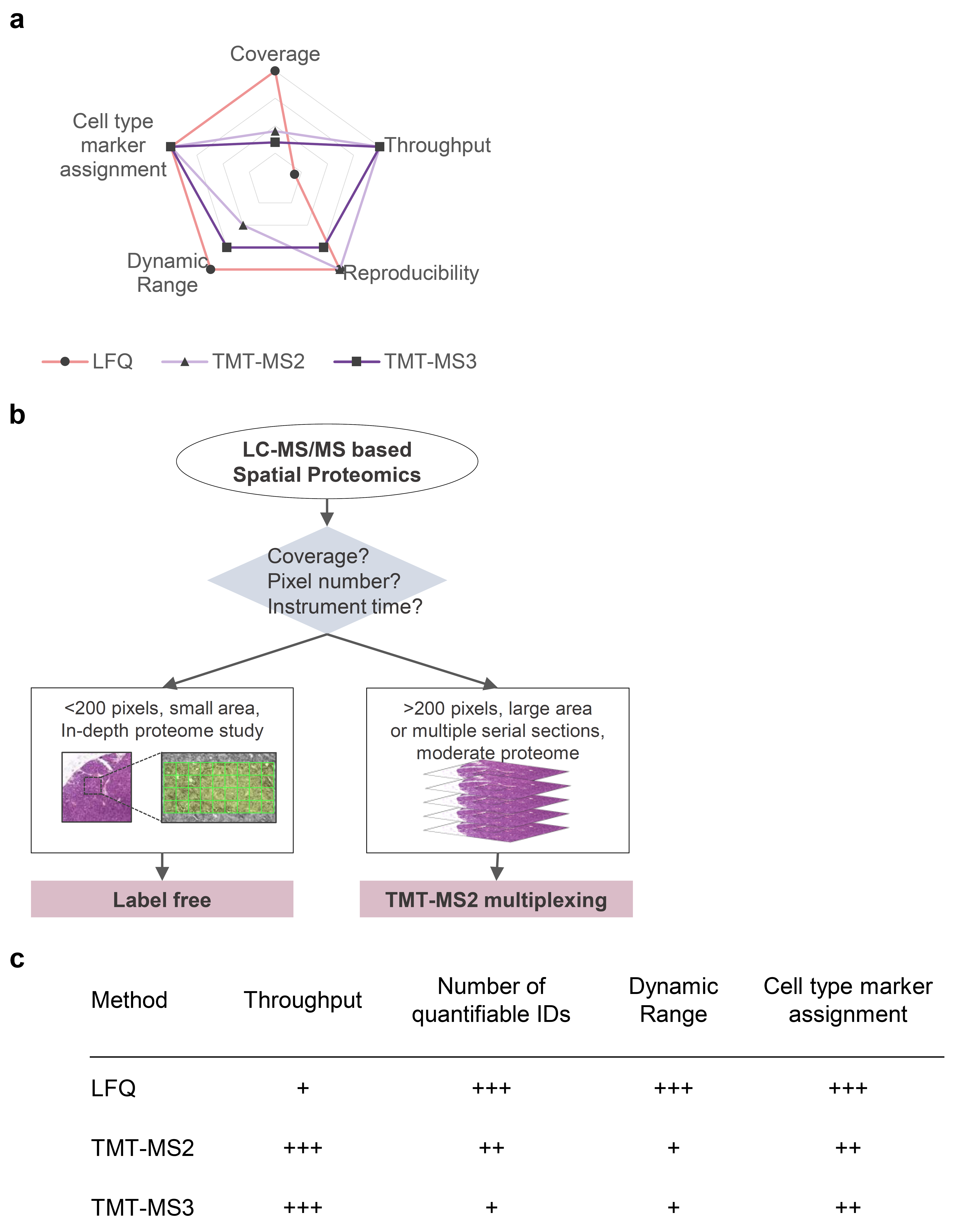

**Supplementary Figure 14. Summary of quantification methods for high-resolution spatial proteomics study.** (**a**) Radar chart for relative comparison of three quantification approaches, including label-free quantification (LFQ), TMT-MS2, and TMT-MS3 analysis based on coverage, sample throughput, reproducibility, dynamic range, and performance for cell-type marker assignment. From the inside to the outside, the outer point represents better performance for a given trait. (**b**) Flow chart for making a decision of the most suitable quantification method for high-resolution proteome mapping based on experimental design. (**c**) A summary table showing the performance of the three quantification approaches.
